# Global analysis of protein-RNA interactions in SARS-CoV-2 infected cells reveals key regulators of infection

**DOI:** 10.1101/2020.11.25.398008

**Authors:** Wael Kamel, Marko Noerenberg, Berati Cerikan, Honglin Chen, Aino I. Järvelin, Mohamed Kammoun, Jeff Lee, Ni Shuai, Manuel Garcia-Moreno, Anna Andrejeva, Michael J. Deery, Christopher J. Neufeldt, Mirko Cortese, Michael L. Knight, Kathryn S. Lilley, Javier Martinez, Ilan Davis, Ralf Bartenschlager, Shabaz Mohammed, Alfredo Castello

**Author notes:** These authors contributed equally to this work (Wael Kamel, Marko Noerenberg, Berati Cerikan).

## Abstract

Severe acute respiratory syndrome coronavirus 2 (SARS-CoV-2) causes COVID-19. SARS-CoV-2 relies on cellular RNA-binding proteins (RBPs) to replicate and spread, although which RBPs control SARS-CoV-2 infection remains largely unknown. Here, we employ a multi-omic approach to identify systematically and comprehensively which cellular and viral RBPs are involved in SARS-CoV-2 infection. We reveal that the cellular RNA-bound proteome is remodelled upon SARS-CoV-2 infection, having widespread effects on RNA metabolic pathways, non-canonical RBPs and antiviral factors. Moreover, we apply a new method to identify the proteins that directly interact with viral RNA, uncovering dozens of cellular RBPs and six viral proteins. Amongst them, several components of the tRNA ligase complex, which we show regulate SARS-CoV-2 infection. Furthermore, we discover that available drugs targeting host RBPs that interact with SARS-CoV-2 RNA inhibit infection. Collectively, our results uncover a new universe of host-virus interactions with potential for new antiviral therapies against COVID-19.

## INTRODUCTION

Severe acute respiratory syndrome coronavirus 2 (SARS-CoV-2) emerged in Wuhan, China, probably as a consequence of zoonotic transmission originating from bats (Zhou et al., 2020). SARS-CoV-2 is the causative agent of coronavirus disease 2019 (COVID-19), and has become a pandemic with more than 51 million infected individuals and more than 1.2 million deaths worldwide (as of 11/11/2020) (Dong et al., 2020). SARS-CoV-2 belongs to the *Coronaviridae* family, and has a single-stranded, positive-sense RNA genome of approximately 30kb. As is the case with all viruses, SARS-CoV-2 is an intracellular parasite that relies on host cell resources to replicate and spread. Intensive efforts have been undertaken to improve our understanding of SARS-CoV-2 interactions with the host cell, including studies to elucidate the transcriptome and proteome dynamics, posttranslational modifications and protein-protein interactions taking place in the infected cell (Banerjee et al., 2020; Bojkova et al., 2020; Bouhaddou et al., 2020; Gordon et al., 2020; Kim et al., 2020b; Klann et al., 2020; Stukalov et al., 2020).

RNA is central for RNA viruses as it does not only function as a messenger for protein synthesis but also as a genome. However, viral genomes cannot encode all the proteins required to mediate, autonomously, all the steps of their lifecycle. Consequently, viruses hijack cellular RNA-binding proteins (RBPs) to promote viral RNA synthesis, translation, stability and assembly into viral particles, (Dicker et al., 2020; Garcia-Moreno et al., 2018). In response, the host cell employs specialised RBPs to detect viral RNAs and intermediates of replication through the recognition of unusual molecular signatures, including tri-phosphate ends, undermethylated cap and double stranded (ds)RNA (Habjan and Pichlmair, 2015). Viral RNA sensing by antiviral RBPs triggers the cellular antiviral state, which leads to the suppression of viral gene expression and the production of interferons. Cellular RBPs are thus crucial regulators of infection, either promoting or restricting virus infection (Garcia-Moreno et al., 2018; Habjan and Pichlmair, 2015). To improve our knowledge of SARS-CoV-2, it is fundamental to elucidate the interactions that viral RNA establishes with the host cell.

Recently, we employed a comprehensive approach, named comparative RNA interactome capture (cRIC), to discover how the RNA-bound proteome (RBPome) responds to the infection with an RNA virus named Sindbis (SINV) (Garcia-Moreno et al., 2019). Strikingly, the RBPome is pervasively remodelled upon SINV infection, and that these changes appear to be critical for both the viral lifecycle and the cell’s antiviral defences (Garcia-Moreno et al., 2019). Interestingly, most of the RBPs that are stimulated by SINV infection redistribute to the membranous structures when this virus replicates, known as viral replication factories, where they interact with viral RNA (Garcia-Moreno et al., 2019). These results suggest that viral RNA is the epicentre of critical interactions with host cell proteins.

In the last few years, a number of approaches have been developed to identify the cellular proteins that interact with viral RNA employing protein-RNA crosslinking and specific RNA capture (Kim et al., 2020a; LaPointe et al., 2018; Ooi et al., 2019; Phillips et al., 2016; Viktorovskaya et al., 2016). While these studies are important advances towards the understanding of viral ribonucleoproteins (RNPs), the choice of the crosslinking and RNA isolation approach leads to distinct depths and specificities. For example, while formaldehyde is a more efficient crosslinker than ultraviolet light (UV), it also promotes protein-protein crosslinks allowing the capture of indirect interactions through protein-protein bridges (Tayri-Wilk et al., 2020). Despite their pros and cons, these studies discovered from dozens to hundreds of cellular proteins that engage with viral RNA in infected cells, highlighting that the viral RNA is a hub for complex host-virus interactions (Kim et al., 2020a; Knoener et al., 2017; LaPointe et al., 2018; Ooi et al., 2019; Phillips et al., 2016; Viktorovskaya et al., 2016).

In this study, we employ multiple proteome-wide approaches to discover which RBPs are involved in SARS-CoV-2 infection. Using cRIC we uncover that the repertoire of cellular RBPs is widely remodelled upon SARS-CoV-2 infection, affecting approximately three hundred proteins involved in a broad range of RNA metabolic processes, antiviral defences and other pathways. Moreover, we employed a newly developed approach named viral RNA interactome capture (vRIC) to comprehensively identify the cellular and viral proteins that interact with viral RNA. We uncover that SARS-CoV-2 RNPs contain dozens of cellular RBPs and several viral proteins, many of which lack known roles in virus infection. For example, we discover that the viral membrane protein (M) and spike (S) interact with SARS-CoV-2 RNA, suggesting potential roles in viral RNA assembly into viral particles. Strikingly, we show that inhibition of cellular proteins that interact with viral RNA using available compounds impairs SARS-CoV-2 infection. Collectively, our data uncover the landscape of protein-RNA interactions that control SARS-CoV-2 infection, including promising targets for novel antiviral treatments against COVID-19.

## RESULTS AND DISCUSSION

### The cellular RNA-binding proteome globally responds to SARS-CoV-2 infection

Cellular RBPs are fundamental for viruses, as they can promote or supress infection (Garcia-Moreno et al., 2019). To elucidate the landscape of active RBPs in SARS-CoV-2 infected cells, we employed cRIC (Garcia-Moreno et al., 2019). In brief, cRIC employs ‘zero distance’, ultraviolet (UV) protein-RNA crosslinking, followed by denaturing lysis, capture of polyadenylated [poly(A)] RNA with oligo(dT) and quantitative proteomics to identify the complement of RBPs that engages with RNA under different experimental conditions (Garcia-Moreno et al., 2019; Perez-Perri et al., 2020; Sysoev et al., 2016). To determine the optimal time post infection for the cRIC experiments, we performed infection kinetics in epithelial human lung cancer cells (Calu-3), a widely used cell model for respiratory viruses. SARS-CoV-2 RNA and infective particles increase over time and peak at 24 hours post infection (hpi) (Figure 1B-C and S1A). Subsequently cell numbers sharply decrease at 36 and 48 hpi, suggesting widespread cell death (Figure 1D). We thus chose two stages of the viral lifecycle: 1) an early timepoint where viral RNA is exponentially increasing (8 hpi), and 2) a late timepoint where viral RNA and extracellular virions peak (24 hpi), prior to the induction of cell death.

**Figure 1.**
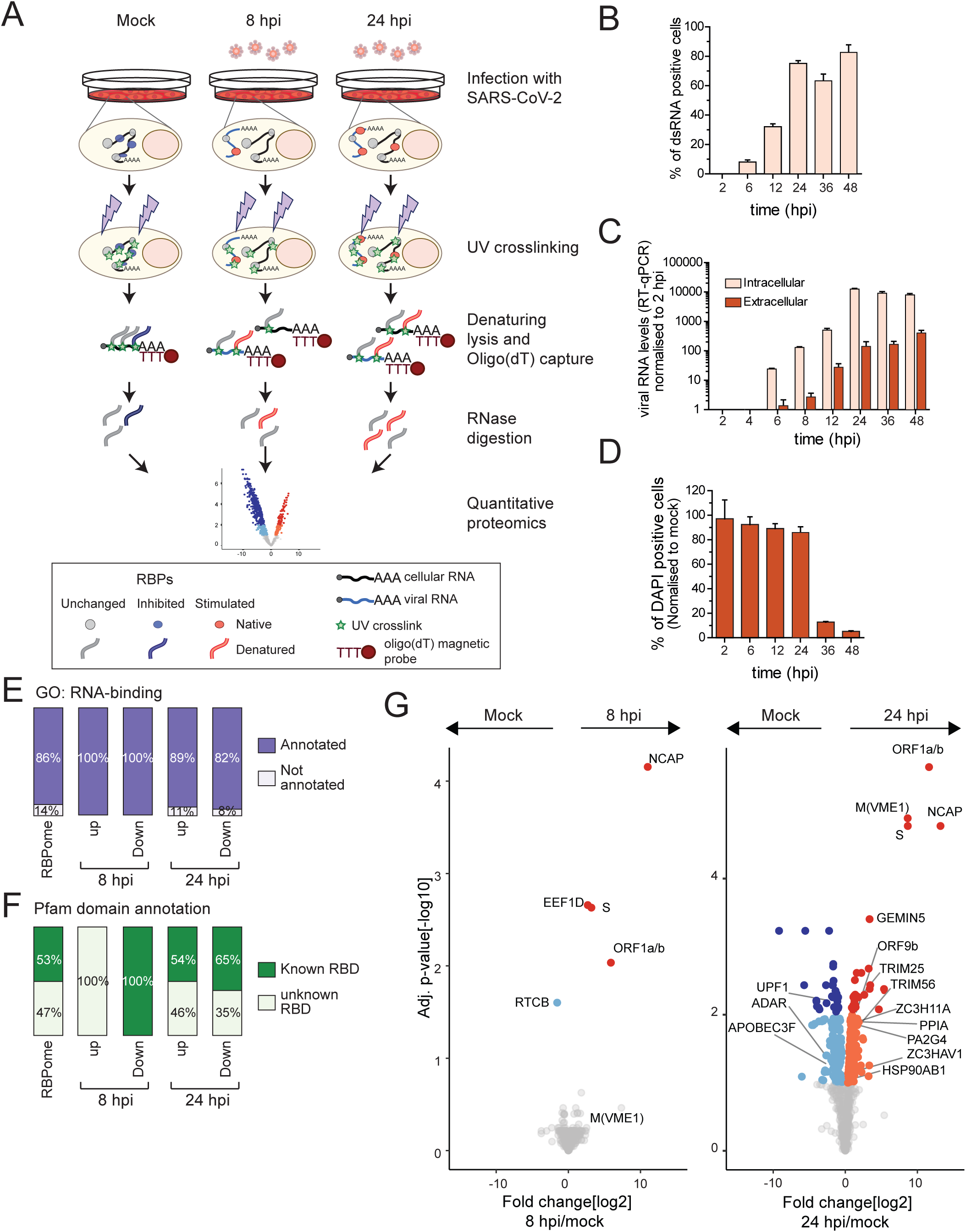
RBPome analysis in SARS-CoV-2 infected cells. A) Schematic representation of cRIC and its application to SARS-CoV-2 infected cells. B) Proportion of infected cells estimated by immunofluorescence using an antibody against dsRNA. C) RT-qPCR analysis of SARS-CoV-2 RNA in cells and isolated from the supernatant of infected cells. Samples were collected at the indicated times post infection. D) Number of adhered cells at different times post SARS-CoV-2 infection counted using DAPI stain. Error bars in B, C and D represent the standard error of the mean (SEM) of three independent experiments. E) Proportion of the identified RBPs that are annotated by the gene ontology (GO) term ‘RNA-binding’. F) Proportion of identified RBPs that harbour known RBDs or lack domains linked to RNA binding. The definition of known RBDs is taken from Garcia-Moreno et al, 2018. G) Volcano plots showing the log2 fold change (x axis) and its significance (adj. p-value, y axis) of each protein (dots) in the cRIC experiments. Fold changes are generated as the ratio between the protein intensity at 8 hpi and in uninfected cells (left) or at 24 hpi and in uninfected cells (right), using data from three biological replicates. Red and dark blue proteins represent RBP upregulated or downregulated, respectively, with 1% false discovery rate (FDR), while orange and cyan represent proteins enriched with 10% FDR.

Once the SARS-CoV-2 infection model was characterised, cRIC was then applied to SARS-CoV-2 infected (8 and 24 hpi) and uninfected cells (Figure 1A). We identified a total of 809 proteins; 86% of which are annotated by the gene ontology term ‘RNA-binding’ and are enriched in well-established RNA-binding domains, resembling previously established *bona fide* RBPomes (Figure 1E-F, S1B and Table S1) (Hentze et al., 2018). 70 proteins displayed changes greater than 2-fold at 8 hpi, although only 5 qualified as statistically significant (Figure 1G and Table S1). This suggests that early RBP responses are either subtle or are variable across replicates. Conversely, 335 RBPs were significantly altered at 24 hpi when compared to uninfected cells. Of these, 176 showed increased and 159 decreased RNA-binding activity (Figure 1G and Table S1). Importantly, SARS-CoV-2-regulation affects both classical RBPs harbouring well-established RBDs and unorthodox RBPs lacking known RBDs (Figure 1F). Moreover, regulated RBPs, and especially those stimulated by SARS-CoV-2, include proteins annotated by GO terms and KEGG pathways related to antiviral response and innate immunity (Figure S1C). Together, these results reveal that SARS-CoV-2 infection initially causes a subtle remodelling of the cellular RBPome (8 hpi) that becomes pervasive by 24hpi, affecting nearly 47% of the detected cellular RBPs. Interestingly, cRIC also identified three viral RBPs at 8 hpi and five at 24 hpi (Figure 1G). These include known viral RBPs such as nucleocapsid (NCAP) and the polyprotein ORF1a/b, as well as proteins previously not known to interact with RNA such as M, S and ORF9b.

### Potential causes for SARS-CoV-2 induced RBPome remodelling

SARS-CoV-2 induces a profound remodelling of the cellular RBPome at 24 hpi. We hypothesised that differential RBP-RNA interactions can simply be a consequence of changes in protein abundance, as previously reported for fruit fly embryo development (Sysoev et al., 2016). To assess this possibility, we analysed the whole cell proteome (WCP) of SARS-CoV-2 infected and uninfected cells (Figure S2A and Table S2). For experimental consistency, we employed the inputs of the cRIC experiments. At 8 hpi, only 69 proteins out of the 4555 quantified exhibited significant changes in abundance, with 29 upregulated and 40 downregulated proteins (Figure 2A, S2B and Table S2). The proportion of differentially expressed proteins increased to 222 at 24 hpi, with 60 upregulated and 162 downregulated proteins (Figure 2A, S2B and Table S2). As expected, all viral proteins increased in abundance as infection progressed (Figure 2A).

**Figure 2.**
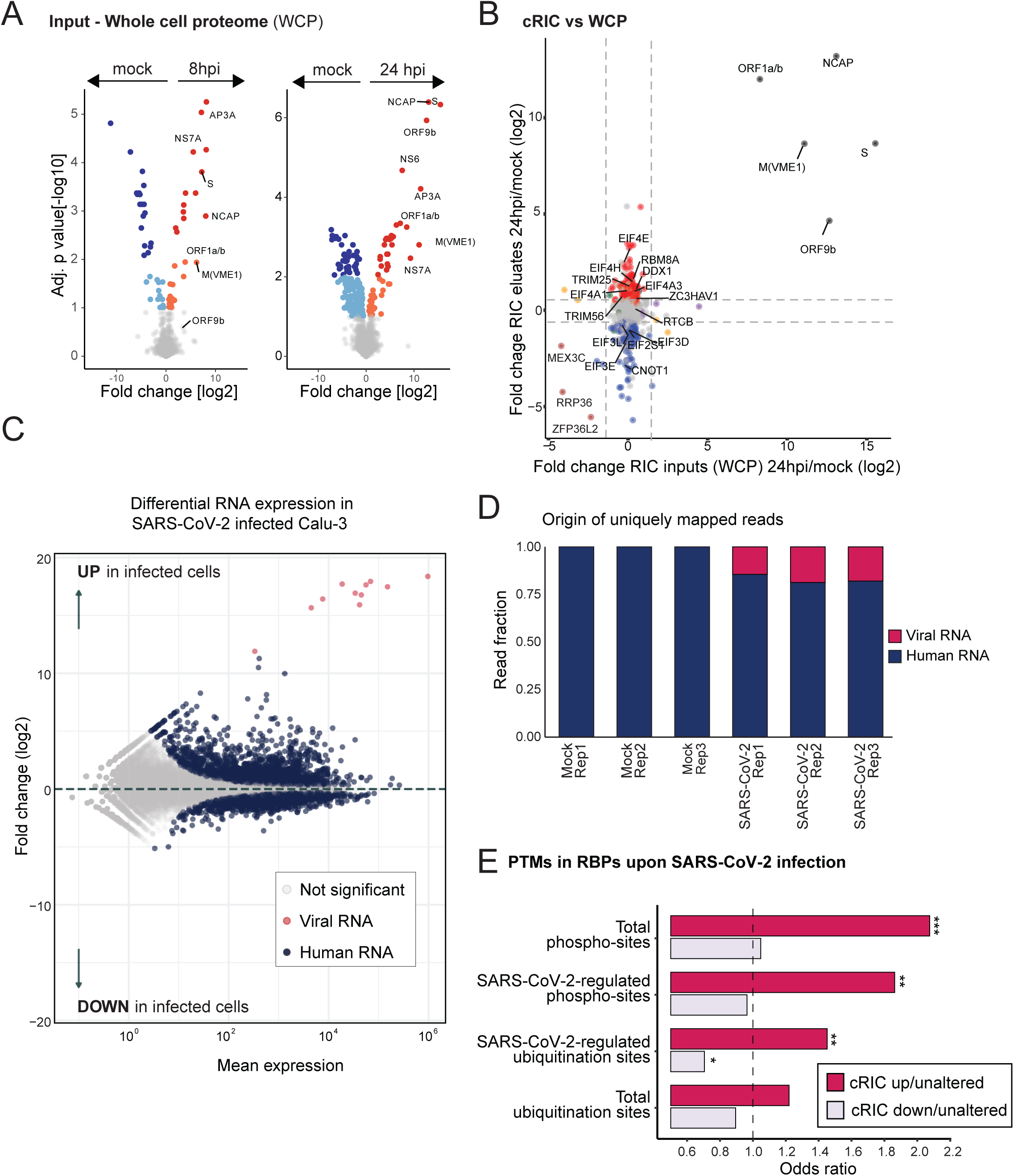
Factors influencing RBP remodelling in SARS-CoV-2 infected cells. A) Proteomic analysis of the inputs of the cRIC experiment corresponding to the whole cell proteome. Volcano plots showing the log2 fold change (x axis) and its significance (adj. p-value, y axis) of each protein between 8 hpi and uninfected conditions (left) or 24 hpi and uninfected conditions (right) using data from three biological replicates. Red and dark blue proteins represent RBP upregulated or downregulated with 1% FDR, while orange and cyan represent proteins enriched with 10% FDR. B) Scatter plot showing the fold changes between 24 hpi and uninfected cell in cRIC (y axis) and the WCP (x axis) for corresponding proteins. In red and blue are the RBPs upregulated or downregulated in the cRIC experiment (FDR<10%) (Figure 1G). C) MA plot highlighting significant changes in gene expression in SARS-CoV-2 infected Calu-3 cells as detected by RNA sequencing. 2733 and 2732 genes are up- and down-regulated, respectively (p-adjusted < 0.01). D) Fraction of uniquely aligned RNA-seq reads mapping to human chromosomes or SARS-CoV-2 genomic RNA in uninfected and infected cells. E) Bar plot showing the odds ratio of previously reported total and SARS-CoV-2 differentially regulated posttranslational modifications (PTMs) in upregulated and downregulated RBPs, relative to the non-regulated RBPs within the cRIC experiment. *, p<0.1; **, p<0.05; ***, p<0.01.

The WCP analysis covers 82% of the proteins identified by cRIC, providing a broad overview on RBP levels in infected and uninfected cells. When we compared cRIC and WCP fold changes, we observed correlation only for viral proteins and a few cellular proteins, which distributed along the diagonal in the scatter plot (Figure 2B). This reflects that the association of viral proteins with RNA increases as viral proteins accumulate. Conversely, changes in cRIC were not matched with similar changes in WCP for most RBPs (Figure 2B). Lack of correlation unequivocally indicates that protein abundance is not a global contributing factor to RBP responses in SARS-CoV-2 infected cells.

RNA abundance can also influence the RBPome, as RNA substrate availability can alter protein-RNA interactions. To test this, we analysed poly(A)-selected RNA sequencing data from Calu-3 cells infected with SARS-CoV-2 for 24h (Blanco-Melo et al., 2020). As expected, SARS-CoV-2 causes substantial alterations in the cellular transcriptome, with 5465 RNAs displaying significant fold changes when compared to the uninfected control (2733 upregulated and 2732 downregulated with p<0.01; Figure 2C and S2C). Particularly, viral RNAs emerge as dominant poly(A) RNA species in the cell, representing 14-19% of the reads (Figure 2C and D). These results have two major implications: 1) that viral RNAs become new abundant substrates for cellular RBPs and 2) that they are captured by oligo(dT) and must thus contribute to the changes observed by cRIC. Therefore, alterations in cellular mRNA abundance and emergence of the viral RNA are likely contributors to the remodelling of the RBPome in SARS-CoV-2 infected cells.

SARS-CoV-2 proteins are known to affect the dynamics and function of cellular RNPs. For example, the non-structural protein (NSP)16 binds the pre-catalytic spliceosomal complex and hampers the splicing reaction, while and NSP1 interacts with the ribosome and inhibits translation initiation on cellular mRNAs (Banerjee et al., 2020; Schubert et al., 2020; Thoms et al., 2020). Hence, transcriptome effects in RBP dynamics can be exacerbated by the alterations that viral proteins induce in the metabolism of cellular RNA.

Post-translational modifications (PTMs) are enriched in protein-RNA interfaces and are known to regulate RBPs (Arif et al., 2018; Castello et al., 2016). We hypothesise that SARS-CoV-2 induced PTMs can thus also affect RBP dynamics. To test this possibility we collected total and SARS-CoV-2 regulated PTMs from available datasets (Bouhaddou et al., 2020; Klann et al., 2020; Stukalov et al., 2020) and mapped them to cRIC-identified RBPs. We found that RBPs, on average, possess considerably more PTMs than other cellular proteins. Of the 335 RBPs regulated by SARS-CoV-2, 123 possessed differential phosphorylation sites and 62 differential ubiquitination sites (Table S3). Strikingly, these SARS-CoV-2-regulated PTMs occur more frequently in upregulated RBPs than in downregulated or unaltered RBPs (Figure 2E), suggesting that PTMs could contribute to the RBP’s ability to interact with RNA. Indeed, we observed that SARS-CoV-2 modulated RBPs were more frequently phosphorylated at multiple sites than their unaltered counterparts (Figure S2E). These results suggest that posttranslational control may also contribute to the differential RNA-binding activity observed for dozens of RBPs in response to SARS-CoV-2 infection.

In summary, the combination of the changes in the transcriptome (Figure 2C-D), the interference of viral proteins with RNA metabolism (Banerjee et al., 2020), and post-translational regulation (Figure 2E and Table S3) are likely contributing to the regulation of RBP activities reported here.

### Kinetics of RBP alterations upon SARS-CoV-2 infection

The kinetics of RBP activation and inhibition can be informative for protein complex dynamics and function. To further characterise the dynamics of RBP after SARS-CoV-2 infection, we clustered proteins based on their cRIC fold changes at 8 and 24 hpi. Our analysis distinguishes eight RBP response profiles as described in Figure 3A and Table S4. Clusters 2 and 7 are dominant, with 114 proteins in each group, reflecting that most RBPs changes are only detected at 24 hpi. By contrast, 70 RBPs exhibited more complex RNA-binding patterns, distributing across clusters 1, 3, 4, 5, 6 and 8.

**Figure 3.**
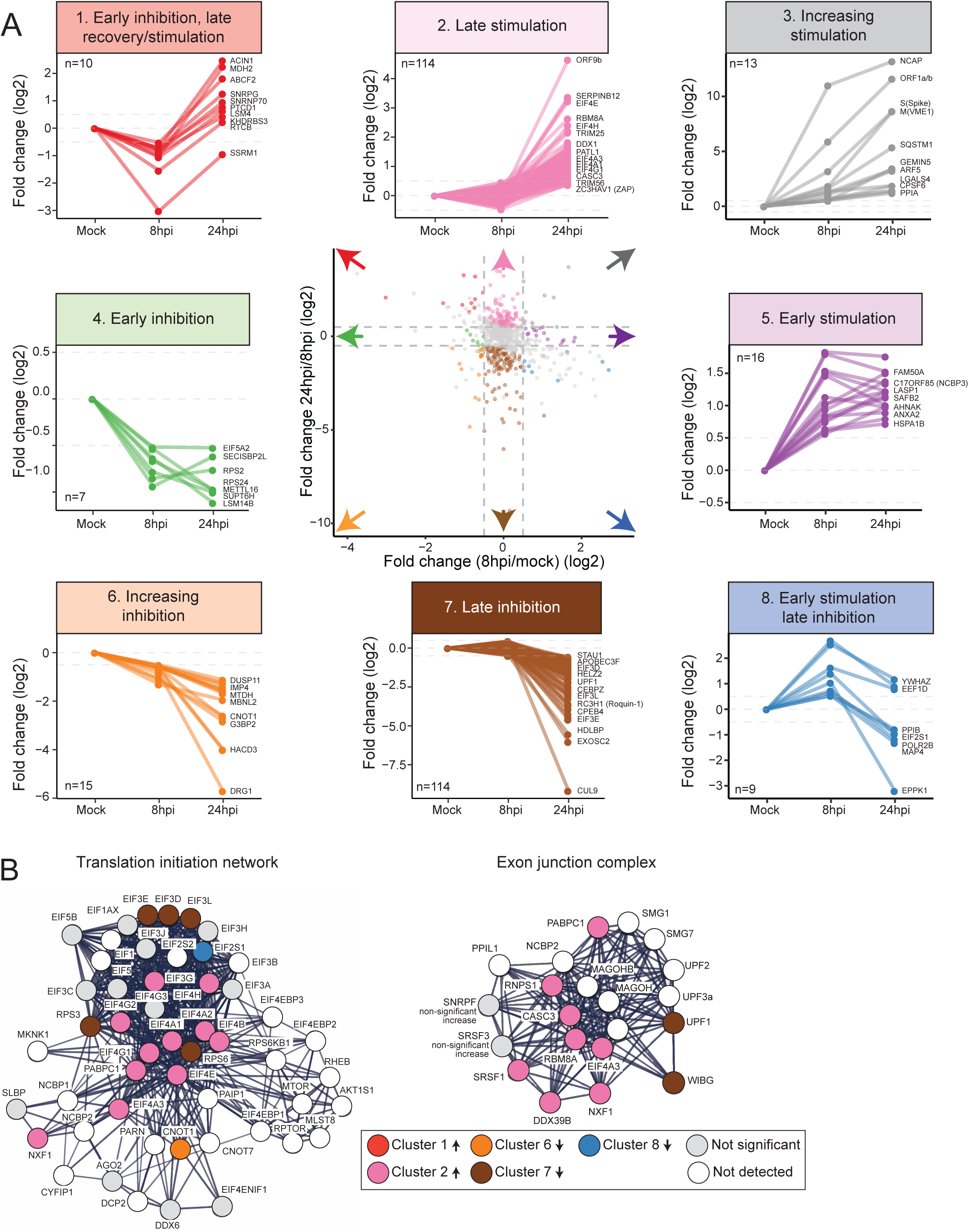
Clustering of RBP responses to SARS-CoV-2 infection. A) Cluster analysis of the RBP dynamics using data from uninfected cells, 8 hpi and 24 hpi. B) Protein-protein interaction network of the translation initiation complex and the exon junction complex generated with STRING. Proteins are represented by circles and are coloured based on their grouping with clusters in panel A.

SARS-CoV-2 RNAs accumulate throughout the infection, and proteins involved in viral replication or its suppression may well display similar kinetics. Accordingly, cluster 3 is comprised of RBPs whose RNA-binding activity increases throughout the infection. Apart from most viral RBPs, cluster 3 harbours several notable cellular factors that have either been linked to virus infection or are known to play critical roles in cellular pathways required for viruses. These include the antiviral protein GEMIN5 (Garcia-Moreno et al., 2019; Martinez-Salas et al., 2020), the autophagy factor SQSTM1 (p62) (Horos et al., 2019), the pre-mRNA cleavage and polyadenylation factor CPSF6, a known cofactor of human immunodeficiency virus type 1 (HIV-1) (Hilditch and Towers, 2014), and the master regulator of virus infection PPIA (cyclophilin A) (Dawar et al., 2017).

SQSTM1 (also p62) is a critical component of the autophagy pathway that plays a key role as a receptor of the autophagy substrates and mediates the interaction with growing phagophores to form autophagosomes (Buscher et al., 2020). In a recent report, it was shown that SQSTM1 autophagy receptor activity is blocked by interaction with vault (vt) RNA 1-1 (Horos et al., 2019). The interaction of SQSTM1 with RNA is mediated by its ZZ and PB1 domain, and the resulting complex is unable to mediate autophagy. The strong increase in RNA-binding activity of SQSTM1 upon SARS-CoV-2 infection suggests that autophagy is inhibited by SARS-CoV-2 through this pathway. Interestingly, the vault complex, which contains vtRNAs, has been reported to reside in close proximity to double-membrane vesicles, which are the sites of viral RNA replication (Klein et al., 2020). However, whether the increase in SQSTM1 RNA-binding activity is mediated by vtRNA1-1, other cellular RNA, or the viral RNA itself requires further investigation.

SARS-CoV-2 NSP1 inhibits protein synthesis by interacting with the ribosome’s mRNA channel (Banerjee et al., 2020; Schubert et al., 2020; Thoms et al., 2020). To determine how this inhibitory interaction affects cellular RBPs, we analysed the kinetic profiles of all proteins annotated by gene ontology (GO) terms related to ‘translation’ and ‘ribosome’. We observed the presence of several components of the eukaryotic initiation factor (EIF)3, EIF2S1 (also EIF2α), elongation factors and ribosomal proteins in clusters 4, 6, 7 and 8, which are comprised of downregulated RBPs (Figure 3A and B, S3A and B and Table S4). Conversely, the cap- and poly(A)-binding proteins eIF4E and PABPC1, as well as the other translation initiation factors such as EIF4A1 and EIF4A2, EIF4B and EIF4G1 and EIF4G3, are present in cluster 2, which is comprised of upregulated RBPs (Figure 3A and B, and Table S4). These opposed results support a model in which the cap- and poly(A)-binding initiation factors can interact with cellular mRNAs but cannot associate with EIF3 and the ribosomal subunit 40S, which agrees with the reported action of NSP1 preventing 40S recruitment to cellular mRNAs (Gehring et al., 2009; Yi et al., 2020).

If this model is correct, it is expected that disrupted translation initiation would increase the presence of the exon junction complex (EJC) onto cellular mRNAs, as it is removed during the pioneering round of translation (Gehring et al., 2009; Yi et al., 2020). To test this hypotheses, we searched for the core components of the EJC in our dataset and observed that EIF4A3, RBM8A and CASC3 exhibit a higher association with RNA in SARS-CoV-2 infected cells (also in cluster 2, Figure 3A and B and S3E). Conversely, the EJC removal factor WIBG (PYM1) (Gehring et al., 2009) is downregulated, further supporting that co-translational removal of EJCs is impaired in infected cells. Moreover, the crucial nonsense mediated decay factor UPF1 (cluster 7) is also inhibited upon infection, which reflects that co-translational quality control is not taking place efficiently. Collectively, these results indicate that SARS-CoV-2 induced protein synthesis shut off may cause the accumulation of matured transcripts into a translation-inactive state.

Deposition of EJCs on cellular RNAs is a consequence of the splicing reaction (Yi et al., 2020). However, a recent study reported that SARS-CoV-2 NSP16 interacts with the U1 and U2 small nuclear (sn)RNAs and disrupt splicing (Banerjee et al., 2020). To assess the effects of NSP16 in RBP dynamics we examined the cRIC fold changes of all spliceosome-associated proteins. Surprisingly, the components of the core spliceosomal complexes showed no significant changes in RNA-binding activity, except for SNRPG that was substantially upregulated (Figure S3D and Table S1). Conversely, several splicing factors showed strong changes in RNA-binding activity, including the branch point binding U2AF2, U2SURP, most serine/arginine (SR)-rich splicing factors (SRSF) and several HNRNPs (Figure S3D, E and F). Many of these proteins play important roles in exon and intron definition and in the recruitment of the spliceosome (Ule and Blencowe, 2019), suggesting potential effects in alternative splicing and splicing efficiency.

### Comparison of SARS-CoV-2 and SINV induced alterations of the RBPome

To determine whether the changes that SARS-CoV-2 induces in the cellular RBPome are shared with other viruses, we compared the SARS-CoV-2 cRIC data to that of SINV (Garcia-Moreno et al., 2019). SINV is a positive stranded virus from the alphavirus genus. As SARS-CoV-2, SINV genome is capped and polyadenylated, although it is substantially smaller (∼11kb vs ∼30kb). Moreover, both viruses produce subgenomic RNAs and replicate in the cytoplasm (Banerjee et al., 2020; Garcia-Moreno et al., 2019). Strikingly, nearly 40% of the changes in RBP activity observed in SARS-CoV-2 were also present in the SINV cRIC dataset (Figure 4A-C). This exciting result indicates that even if these viruses belong to different families and have little or no sequence homology, they cause similar alterations in the RBPome. These consistent changes were observed in both upregulated and downregulated RBPs (Figure 4A and B). Several antiviral factors were noticeable amongst the 93 RBPs showing consistent responses to both viruses, TRIM25, TRIM56, ZC3HAV1 (also ZAP), DHX36 and GEMIN5 (Figure 4D and S4A). These antiviral RBPs are upregulated in both datasets, suggesting that they are likely involved in the antiviral response against both SARS-CoV-2 and SINV (Figure 4D and S4A). TRIM25 is an E3 ubiquitin ligase whose catalytic activity is triggered by RNA binding (Choudhury et al., 2017). It interacts with SINV RNA and has a powerful antiviral effect in infection (Garcia-Moreno et al., 2019). TRIM25 antiviral activity is thought to be mediated by the ubiquitination of RIGI and ZC3HAV1/ZAP (Gack et al., 2007; Li et al., 2017). While RIGI was not detected in our analysis, ZC3HAV1/ZAP RNA-binding activity was upregulated in response to infection (Figure 4D). This suggests that ZC3HAV1/ZAP is potentially the effector TRIM25 antiviral function in cells infected with SARS-CoV-2 and SINV. Another E3 ubiquitin ligase, TRIM56, displayed enhanced RNA-binding activity in both SARS-CoV-2 and SINV infected cells. We recently described that TRIM56 interacts with SINV RNA and supresses viral infection (Garcia-Moreno et al., 2019), and our present data indicates that this may also occur in SARS-CoV-2 infected cells. GEMIN5 is an antiviral factor that interacts with the cap and 5’UTR of SINV RNA and supresses viral mRNA translation (Garcia-Moreno et al., 2019; Martinez-Salas et al., 2020). Given that SARS-CoV-2 RNAs are also capped, it is thus plausible that GEMIN5 hampers SARS-CoV-2 gene expression following a similar mechanism. Other RBPs with prominent roles in virus infection were consistently upregulated by SARS-CoV-2 and SINV, including PPIA (cyclophilin A), PA2G4, ZC3H11A, DDX3, and HSP90AB1 (Figure S4B) (Dawar et al., 2017; Garcia-Moreno et al., 2019; Valiente-Echeverria et al., 2015; Younis et al., 2018).

**Figure 4.**
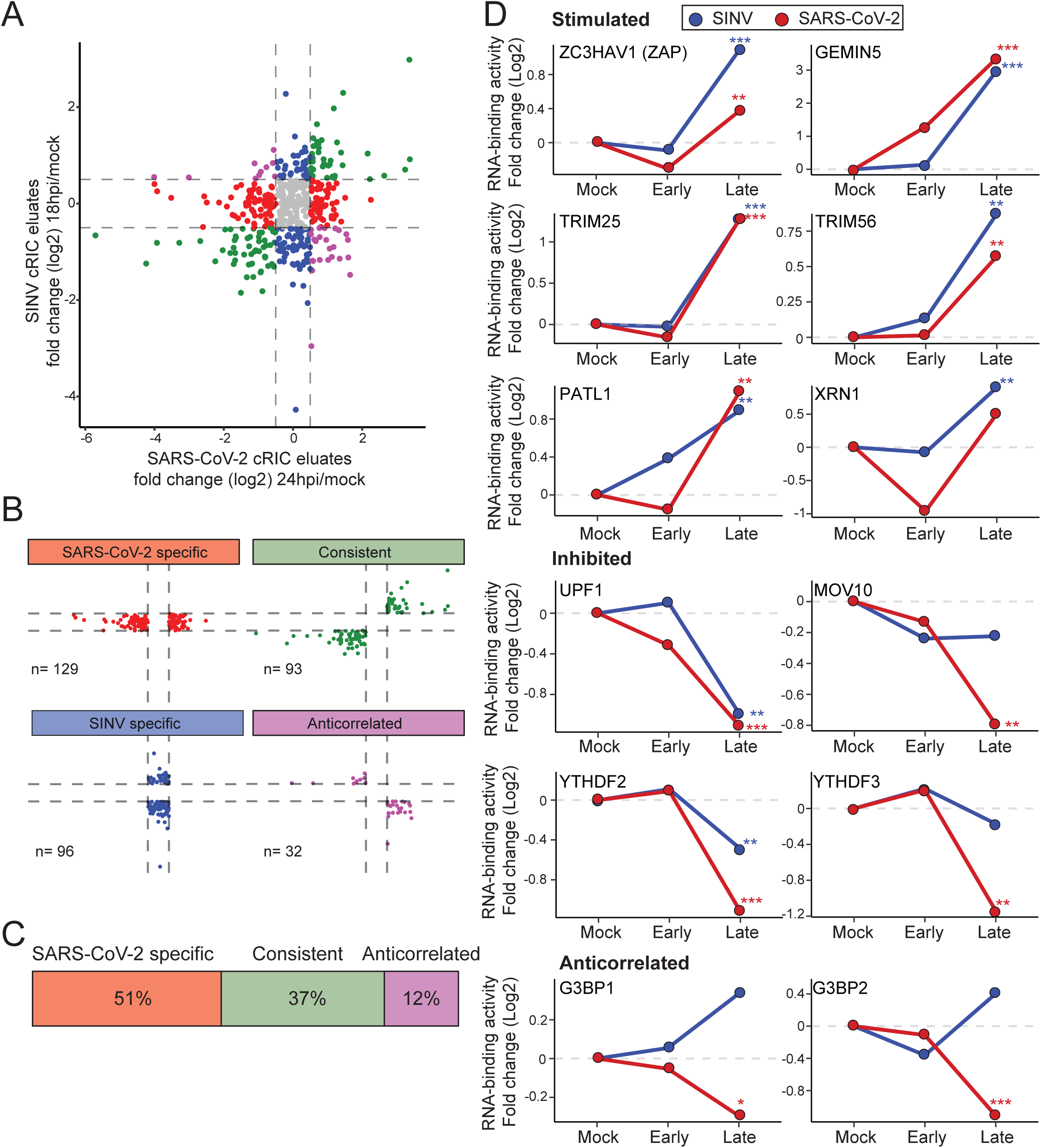
Analysis of RBP dynamics in SARS-CoV-2 and SINV infected cells. A) Scatter plot of the fold change between infected and uninfected cells, using the data from the cRIC experiments in SARS-CoV-2 infected cells (Figure 1G) and in SINV-infected cells (Garcia-Moreno et al., 2019). We used the late time post infection in both experiments, which corresponds to 24 hpi for SARS-CoV-2 and 18 hpi for SINV. Proteins were grouped based on their behaviour (B) and the overlap between datasets was estimated (C). D) Fold change of selected proteins in SARS-CoV-2 (red) and SINV (blue) cRIC analyses. ‘Early’ represents 8 hpi for SARS-CoV-2 and 4 hpi for SINV. ‘Late’ is 24 hpi for SARS-CoV-2 and 18 hpi for SINV *, FDR < 20%; **, FDR < 10% and *** FDR < 1%.

Our data also revealed antiviral RBPs that are downregulated by SARS-CoV-2 and, in several instances, also by SINV. These include the RNA editing enzymes ADAR, APOBEC3F and APOBEC3G, and the nonsense mediated decay helicase UPF1 (Figure S4C).

Interestingly, 12% of the proteins exhibited opposite behaviour in the two viral models. Many of these can be traced back to membraneless organelles such us the paraspeckles and stress granules. The core paraspeckle components NONO, PSPC1, SFPQ and MATR3 display opposite trends, being repressed by SINV and stimulated or unaffected by SARS-CoV-2 (Figure S4D). It is proposed that paraspeckles are critical to sequester proteins and/or mRNAs to regulate gene expression, although the importance of paraspeckle proteins in virus infection remains poorly understood and deserve further consideration (Fox et al., 2018). Similar anticorrelation was observed with the stress granule proteins G3BP1 and G3BP2 (Figure 4D). Stress granules are protein-RNA assemblies that play a defensive role against viruses by sequestering viral RNA (McCormick and Khaperskyy, 2017). Alphaviruses like SINV are known to supress stress granule formation, and this is accompanied by an increase of G3BP1 and G3BP2 RNA-binding activity (Garcia-Moreno et al., 2019; Kim et al., 2016; Panas et al., 2012; Scholte et al., 2015). The inhibition of G3BP1 and G3BP2 in SARS-CoV-2 infected cells may thus reflect an opposite outcome, i.e. lower association with RNA due to the induction of stress granules.

### The SARS-CoV-2 RNA interactome

The cRIC method captures both SARS-CoV-2 and cellular mRNAs, which represent 14-19% and 84-80% of the eluted RNA, respectively (Figure 2D and S2D). Therefore, it is not possible to know *a priori* which of the observed protein-RNA interactions are driven by viral RNA. To systematically identify the RBPs that interact directly with SARS-CoV-2 RNAs, we applied a newly developed approach that we named viral RNA interactome capture (vRIC) (Figure 5A and B and S5A) (Kamel et al., In preparation). In brief, SARS-CoV-2-infected and uninfected Calu-3 cells are treated with the RNA polymerase II (RNAPII) specific inhibitor flavopiridol (Fvo), followed by a pulse with the photoactivatable nucleotide analogue 4-thiouridine (4SU). As viral RNA polymerases are insensitive to Fvo, temporal inhibition of RNAPII causes 4SU to be predominantly incorporated into nascent viral RNAs. Cells are then UV irradiated at 365 nm to induce crosslinks between viral RNA and proteins placed at a ‘zero distance’ from the 4SU molecules. As natural nucleotide bases do not absorb UV light at 365 nm, protein-RNA crosslinking is restricted to 4SU-containing viral RNA. Cells are then lysed under denaturing conditions and poly(A)-containing RNA is captured with oligo(dT) following a previously designed robust procedure (Castello et al., 2012). After elution, proteins co-purified with the viral RNA are analysed by quantitative label free mass spectrometry.

**Figure 5.**
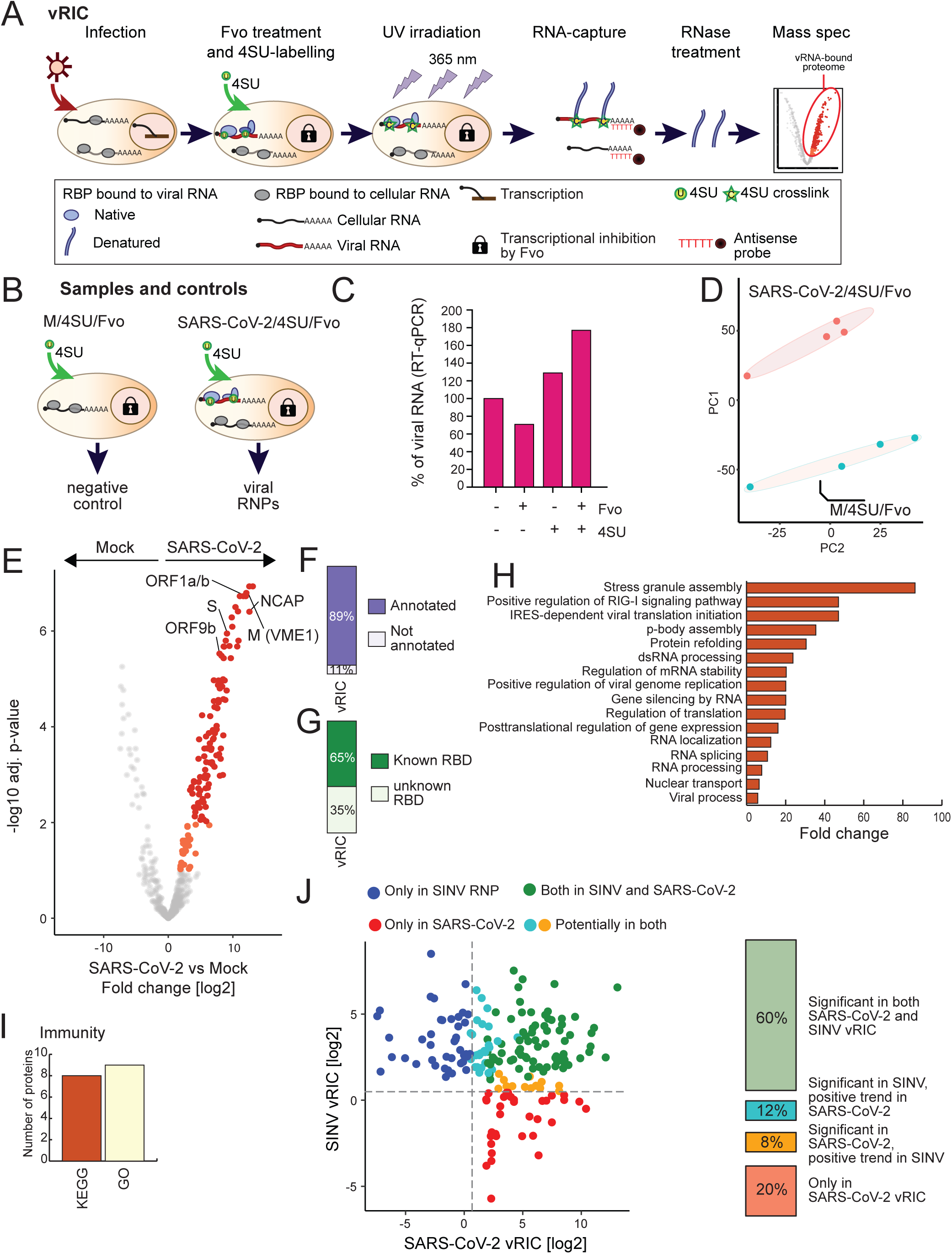
vRIC analysis of the SARS-CoV-2 RNA interactome. A) Schematic representation of vRIC. B) Controls used in the vRIC experiment are schematised (expanded in Figure S5A). C) The effects of 4SU and Fvo on SARS-CoV-2 RNA levels were analysed by RT-qPCR. D) Principal component analysis (PCA) of the four vRIC replicates, including samples from infected (SARS-CoV-2/4SU/Fvo) and mock (M/4SU/Fvo) cells. E) Volcano plots showing the log2 fold change (x axis) and significance (adj. p-value, y axis) of each protein in the vRIC experiment. Fold changes are estimated as the ratio between the protein intensity in eluates from infected (SARS-CoV-2/4SU/Fvo) and mock (M/4SU/Fvo) samples using data from four biological replicates. Red dots represent proteins enriched in samples from SARS-CoV-2 infected cells at 1% FDR, while orange represents proteins enriched with 10% FDR. F) Proportion of the proteins enriched in the SARS-CoV-2 RNPs that are annotated as ‘RNA-binding’ in gene ontology (GO). G) Proportion of the proteins enriched in the SARS-CoV-2 RNPs that harbour known RBDs or lack domains linked to RNA binding. H) GO enrichment analysis of the proteins enriched in the SARS-CoV-2 RNPs. I) Proportion of the proteins enriched in SARS-CoV-2 RNPs that are annotated to immunity related terms and pathways in GO or KEGG. J) Scatter plot showing the fold change between infected and uninfected cells, using the vRIC data from SARS-CoV-2 infected cells (Figure 1G) and SINV-infected cells (Kamel et al., in preparation). In the right, box plot showing the overlapping between the two datasets.

Our control experiments showed that neither Fvo nor 4SU interfered with SARS-CoV-2 replication under the conditions used (Figure 5C). In mock cells, 4SU incorporation followed by 365nm UV crosslinking and oligo (dT) capture led to the isolation of the steady state RBPome (Figure S5B-F). However, when 4SU was omitted or Fvo was added, the amount of proteins co-isolated with RNA was massively reduced in both silver staining and proteomic analyses (Figure S5B-F). These results show that it is required active RNAPII in uninfected cells to achieve efficient 4SU-dependent protein-RNA UV crosslinking. Conversely, when cells were infected with SARS-CoV-2, efficient protein isolation was observed despite Fvo treatment (Figure 5D-E and S5B-F). These findings confirm that 4SU incorporation into nascent viral RNAs in presence of Fvo promotes effective UV protein-RNA crosslinking at 365nm (Figure 5D-E and S5B-F). In agreement, a principal component analysis revealed that the proteomic datasets derived from uninfected and SARS-CoV-2 infected cells are clearly distinct (Figure 5D), with a total of 139 RBPs enriched in vRIC eluates from SARS-CoV-2 infected cells (SARS-CoV-2/4SU/Fvo) over the mock control (M/4SU/Fvo), 107 with 1% false discovery rate (FDR) and 32 additional proteins with 10% FDR (Figure 5E, Table S5). 89% of the proteins within the SARS-CoV-2 mRNA interactome are annotated with the GO term ‘RNA binding’ and are enriched in known RBDs (Figure 5F-G), supporting the capacity of vRIC to identify *bona fide* protein-RNA interactions. While 65% of the enriched RBPs harbour known RBDs, vRIC was also able to identify unorthodox RBPs with unknown RBDs that interact with SARS-CoV-2 RNAs (Figure 5G and Table S5). The SARS-CoV-2 RNA interactome is enriched in GO terms associated RNA metabolism (RNA splicing, transport, stability, silencing and translation), antiviral response (e.g. RIGI pathway), cytoplasmic granule assembly (stress granules and P-bodies), and virus biology (e.g. viral process, dsRNA binding, IRES-dependent viral RNA translation) (Figure 5H). Notably, 8 and 9 proteins were annotated by innate immunity related terms in KEGG and GO, respectively (Figure 5I).

To determine to what extent the SARS-CoV-2 RNA interactome harbours cellular RBPs that are also present in the RNPs of other viruses, we compared the SARS-CoV-2 vRIC to a SINV vRIC dataset generated in a parallel study (Kamel et al., In preparation). The SARS-CoV-2 vRIC dataset is smaller than the SINV counterpart, likely due to the limited starting material available (Figure S5G). Nevertheless, 60% of the RBPs within the SARS-CoV-2 RNA interactome were also present in that of SINV (Figure 5J). These striking results suggest that viral RNPs may share a larger proportion of cellular factors than previously anticipated, opening the possibility to target commonly used RBPs in broad-spectrum therapeutic approaches.

The cRIC analysis revealed global alterations of the translation machinery, likely due to the NSP1-induced shut-off of cellular mRNA translation (Figure 3B and S3A-B). To test if these alterations also apply SARS-CoV-2 RNAs, we examined the translation factors present in the viral RNP. Most of the proteins involved in the recognition of the cap and poly(A) tail are identified in SARS-CoV-2 RNP, including EIF4G1, EIF4G3, EIF4A1, EIF4A2, EIF4B and PABPC1 (Figure 5E and Table S5). However, one of the critical components is missing: the cap-binding protein EIF4E. While we cannot rule out that this missing protein is a false negative, other capped RNA viruses such as SINV can initiate translation without EIF4E, calling for further experiments to discriminate between these two possibilities (Carrasco et al., 2018). Moreover, several core EIF3 subunits (A, C, D and G) are highly enriched in the SARS-CoV-2 RNP, revealing that the molecular bridge connecting the ribosome and the mRNA via interaction with EIF4G (Merrick and Pavitt, 2018) is active in SARS-CoV-2 mRNAs despite the downregulation of several EIF3 subunits in the cRIC analysis (Figure 5E and Table S5). These results suggest that even though EIF3 subunits C and D have an overall reduced association with RNAs likely due to NSP1 action on translation, they do interact with SARS-CoV-2 RNA to enable viral protein synthesis.

cRIC revealed an upregulation of many HNRNPs (Figure S3F). To test if viral RNA is involved in these alterations, we examined the vRIC dataset. Notably, 10 HNRNPs interact with SARS-CoV-2 RNA, particularly from the A family (A0, A1, A2B1, A3, C, DL, M, L, Q [SYNCRIP] and R). Indeed, immunofluorescence analysis revealed that a subpopulation of HNRNPA1 accumulates in the viral replication factories (labelled with anti-dsRNA), confirming these results (Figure 5SH). These results suggest that the enhancement of HRNP RNA-binding activity may be driven by SARS-CoV-2 RNA accumulation.

The cRIC analysis revealed a connection between SARS-CoV-2 infection and RNA granules (Figure 4D and S4D). To determine if such interplay involves the viral RNA, we searched for known components of RNA granules in the vRIC dataset. Notably, we noticed the presence of core stress granule components G3BP1 and G3BP2, and their interacting proteins CAPRIN1, NUFIP2 and USP10 within SARS-CoV-2 RNPs (Figure 5E and Table S5). These results, together with the observed downregulation of G3BP1 and G3BP2 RNA-binding activities (Figure 4D and S4E) and the known interaction between G3BP1 and the viral nucleocapsid (NCAP) (Gordon et al., 2020), reflect an intimate relationship between stress granules and SARS-CoV-2 RNAs. Additionally, the P-body components DDX6, LSM14A, PATL1 and the miRNA mediator AGO2, also interact with SARS-CoV-2 mRNA. Conversely, none of the nuclear paraspeckle proteins were statistically enriched in the viral RNP, suggesting that their role in SARS-CoV-2 infection, if any, might be indirect. Collectively, our data shows that SARS-CoV-2 RNA engage with components of stress granules and P-bodies. The biological relevance of these interactions deserves further characterisation.

SARS-CoV-2 RNA is post-transcriptionally edited, although the importance of this remains unknown (Kim et al., 2020b). To get more insights into this phenomenon and its consequences in the composition of the viral RNP, we searched for all ‘editors’ and ‘readers’ that interact with SARS-CoV-2 RNAs (Table S5). ADAR is downregulated upon SINV infection (Table S1); however, it is highly enriched in SARS-CoV-2 RNPs (Figure 5E and Table S5). It catalysed the conversion of adenosines to inosine, which can affect several aspects of RNA function, including structure, RBP binding sites and coding sequence, potentially hampering viral replication. The participation of ADAR in SARS-CoV-2 infection is underscored by a recent study reporting adenosine deamination in the SARS-CoV-2 RNA (Di Giorgio et al., 2020). Methyl 6 adenosine (m6A) also plays critical roles in virus infection and viral RNA is typically enriched with this modification (Tan and Gao, 2018). This molecular signature is recognised by a family of proteins known as ‘readers’, which regulate RNA fate (Wang et al., 2014). While the readers YTHDF2 and YTHDF3 are downregulated in the cRIC analyses of both SINV and SARS-CoV-2 infected cells, YTHDC1 and YTHDC2 are stimulated (Figure 4D and S4F and G). These opposed results indicate that YTH m6A readers are differentially regulated in response to infection. Our vRIC analysis shows that YTHDC2 is significantly enriched in the SARS-CoV-2 RNPs (Figure 5E, Table S5). Taken together, these results support the potential role of YTHDC2, and perhaps YTHDC1, as mediators of m6A function in SARS-CoV-2 infection.

Other noteworthy proteins that interact with SARS-CoV-2 RNA include five helicases (DDX1, DDX3X, DDX6, DDX60, DHX57); four chaperones (HSP90AA1, HSP90AB1, HSPA5 and HSPA8); the actin-interacting proteins SYNE1 and SYNE2; the E3 ubiquitin ligase MKRN2; the vesicle membrane protein VAT1 that interacts with M, ORF7b (NS7B) and ORF9b (Gordon et al., 2020); the glycolysis enzyme PKM that is known to moonlight as an RBP (Castello et al., 2012); the antiviral protein OASL that belongs to a family of SARS-CoV-2 susceptibility factors (Pairo-Castineira et al., 2020), and three separate subunits of the protein phosphatase 1 (PPP1CA, PPP1CB, PPP1R3A), amongst others. Collectively, vRIC shows that SARS-CoV-2 RNA engages with a broad range of cellular RBPs, including classical and non-canonical RNA binders. This dataset represents a step forward towards understanding the roles of cellular proteins in the function of SARS-CoV-2 RNPs.

### Viral proteins that interact with viral and cellular RNA

The cRIC and vRIC data are in full agreement regarding the SARS-CoV-2 proteins that interact with RNA, even though these two methods employ different crosslinking chemistries (Castello et al., 2012). Viral RBPs include the polyprotein ORF1a/b, NCAP, and, surprisingly, M, S and ORF9b (Figure 1E-F, 5E, 6A-B and S6A). To determine which type of interaction these proteins establish with RNA, we normalised the protein intensity in vRIC and cRIC by that of the WCP (Figure 6A-B, S6A and Table S6). NCAP and ORF1a/b displayed the highest UV ‘crosslink-ability’, followed by M, ORF9b and S. Generally, the efficiency of protein-RNA UV crosslinking depends on several factors, including 1) the geometry of the protein-RNA interaction and, in particular, the quantity and quality of the contacts to the nucleotide bases, 2) the physicochemical properties of the bases and amino acids in close proximity, 3) the duration of the interaction, and 4) the proportion of the protein that engages in RNA binding. Taking these factors into consideration, we can suggest that ORF1a/b and NCAP establish optimal and stable interactions with RNA, while M, ORF9b and, especially, S mediate shorter-lived and/or geometrically less favourable interactions for crosslinking (e.g. interactions with the phosphate backbone). However, the high protein sequence coverage and peptide intensity in both vRIC and cRIC experiments (Figure 6C-E and S6B) strongly support the possibility that these protein-RNA interactions are not false positives.

**Figure 6.**
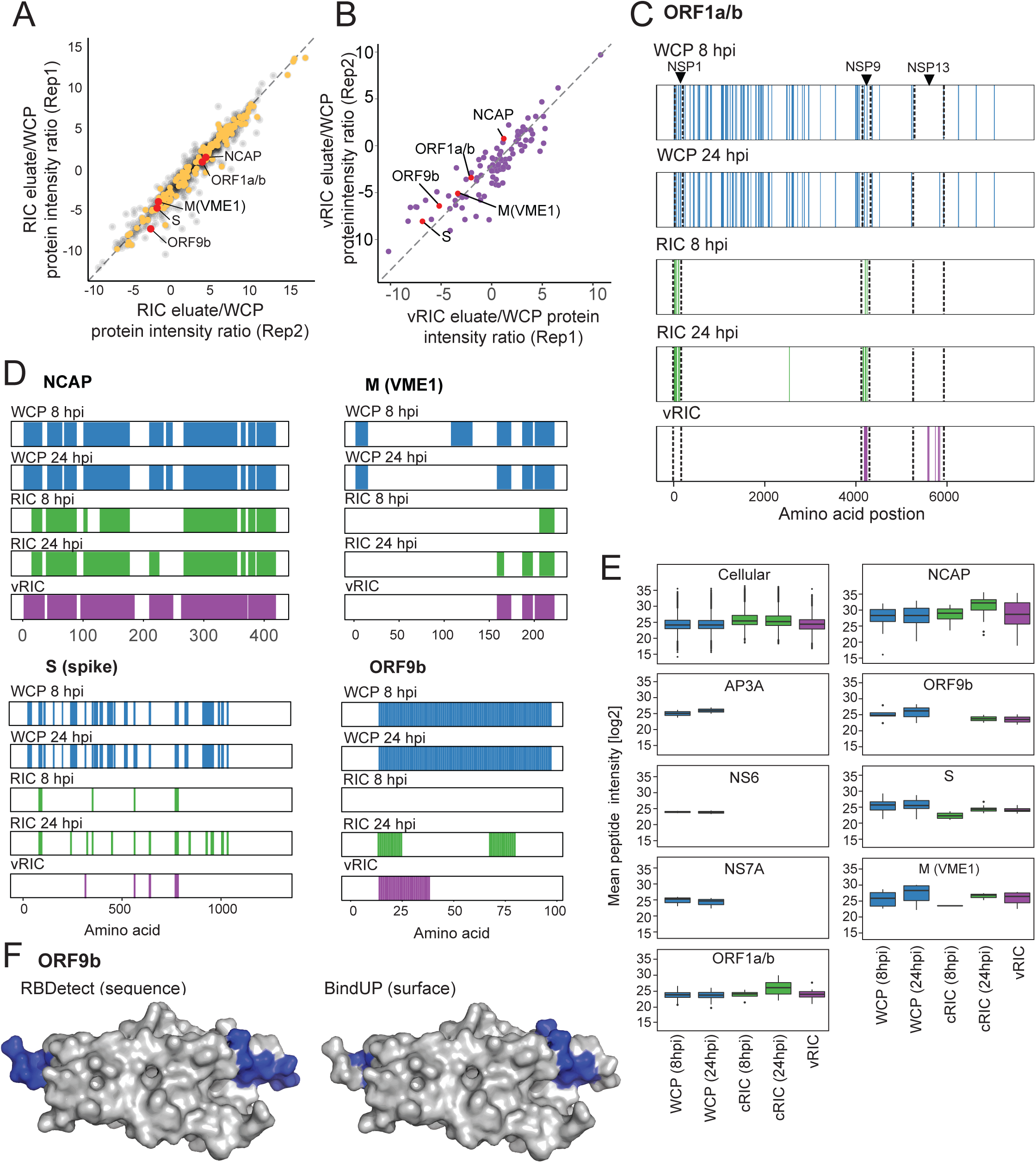
SARS-CoV-2 proteins that interact with RNA. A and B) Representative scatter plot showing the cRIC (left, 24 hpi/mock)) or vRIC (right, infected/uninfected) fold change normalised to the fold change in the WCP (24 hpi/mock). Replicates 1 and 2 were chosen as illustrative examples and the rest of comparisons can be found in Figure S6A. Cellular RBPs upregulated in the cRIC experiments (Figure 1G) are coloured in yellow, cellular RBPs enriched in SARS-CoV-2 vRIC (Figure 5E) are shown in violet, and viral proteins in red. C and D) Sequence coverage analysis. Peptides detected in WCP (blue), cRIC (green) and vRIC (violet) are mapped to the viral proteins plotted from N-terminus to C-terminus (x axis). E) Boxplot showing peptide intensity distribution in cRIC, vRIC and WCP for each of the viral proteins detected. Colours as in C-D. G) ORF9b structure showing the protein surface (PDB ID: 6z4u). Peptides with high probability of RNA binding by RBDetect (left) or BindUP (right) are coloured in blue.

ORF1a/b is a polyprotein comprising of 16 mature polypeptides. While the peptides detected in the WCP mapped uniformly throughout the polyprotein, both cRIC and vRIC identified peptides clustered only in specific regions (Figure 6C). The first peptide cluster mapped to NSP1 and was only detected by cRIC (both 8 and 24 hpi). The lack of signal in vRIC samples strongly indicates that NSP1 does not interact with viral RNA but cellular mRNAs, which are highly enriched in the eluates of the oligo(dT) capture (Figure S2D). It is well established that NSP1 interacts with the ribosome channel to block cellular mRNA translation (Banerjee et al., 2020; Schubert et al., 2020; Thoms et al., 2020). However, how does NSP1 disrupt the translation of cellular mRNAs and not viral RNAs? NSP1’s interaction with cellular mRNAs could contribute to this discrimination. The second peptide cluster mapped to NSP9 and is present in both vRIC and cRIC (Figure 6C). The detection of NSP9 by vRIC agrees well with its known role in viral replication and the well-established interaction of its SARS-CoV-1 orthologue with single stranded RNA (Chandel et al., 2020; Egloff et al., 2004). Whether NSP9 is restricted to viral RNA or can also act on cellular mRNAs requires further characterisation. The third peptide cluster mapped to the RNA helicase NSP13, which is critical for SARS-CoV-2 replication (Chen et al., 2020). Cluster 3 peptides are only detected by vRIC, which supports that NSP13 only interacts with viral RNA. Together, these data reveal that at least seven viral proteins interact with RNA in infected cells.

The proteins M and S also reliably and robustly co-purify with RNA upon cRIC and vRIC, and this is evidenced by several high intensity peptides (Figure 1E-F, 5E, 6A-B, E and S6B). The most likely scenario in which these proteins could engage with viral RNA is during virus assembly and within viral particles (Klein et al., 2020; Yao et al., 2020). To determine if M and S have sequences compatible with RNA binding, we used RBDetect, a software package that employs shrinkage discriminant analysis to predict peptide segments that interact with RNA. Strikingly, we detected two segments in the intravirion region of M that share sequence similarities with *bona fide* RNA-binding sites present in cellular RBPs (Figure S6C), supporting its observed ability to interact with RNA. Similarly, the intravirion part of S also harbours a ∼15 amino acid motif compatible with RNA binding (Figure S6C). Both M and S RNA-binding regions are present in both SARS-CoV-2 and SARS-CoV-1, suggesting that the underlying functional sequences are conserved. While we cannot fully rule out that these interactions with RNA are stochastic due to protein-RNA proximity in the context of the virion, their prominence in the vRIC and cRIC data suggest that they may play a role in infection (Figure 6C-E and S6B). For example, they may contribute to the recruitment of viral RNA, or to the budding and/or structural arrangement of the viral particle. Importantly, a recent report shows that NCAP clusters form underneath the viral envelope during budding of the viral particles, and this structure persists in the mature particles (Klein et al., 2020; Yao et al., 2020). The observed RNA-binding activity of S and M may contribute to recruit and/or trap viral RNA underneath the double membrane at the budding site. Indeed, M is known to recruit NCAP to the budding sites in SARS-CoV-1 infected cells (Ye et al., 2004). In agreement, cryo-electron tomography analysis of infected cells revealed that membrane invagination at the budding site appears to require the presence of NCAP (Klein et al., 2020), implying a potential role for RNA in the process of particle formation.

The viral protein ORF9b was also consistently identified by both cRIC and vRIC, supporting that it is a novel RNA-binding protein (Figure 1E-F, 5E and 6A-B). Very little is known about ORF9b beyond its ability to interfere with interferon responses (Jiang et al., 2020). To determine if ORF9b also contain sequences compatible with RNA binding, we used RBDetect (sequence-based software) and, given the availability of a deposited structure (6Z4U) (Weeks et al., In preparation), we also considered surface physicochemical properties (BindUP) (Paz et al., 2016). Both approaches agree that there is a discrete region in ORF9b that generates a positively charged surface with high probability to interact with nucleic acids (Figure 6G and S6C and D). Further work is required to define the role of the RNA-binding activity of ORF9b in SARS-CoV-2 infection.

Taken together, our data reveal seven viral proteins harbouring RNA-binding activities, highlighting M, S, ORF9b and novel viral RBPs.

### Functional importance of protein-RNA interactions in SARS-CoV-2 infection

To determine if our study has potential for discovery of new regulators of SARS-CoV-2 infection, we assessed the incidence of vRIC and cRIC identified proteins in genome wide screens with other viruses. The superset comprises of studies using RNA interference (RNAi), CRISPR-Cas9, and haploid line screens for 36 viruses, including RNA and DNA viruses (Table S7). This analysis revealed that cRIC and vRIC identified 47 RBPs that that frequently occur in functional screenings (>3 studies), causing phenotypes in virus infection (Figure 7A, B, S7A, Table S7). Moreover, we used an automated PubMed search pipeline to assess how many RBPs have been robustly linked to virus infection in the literature. Interestingly, 73 (43.5%) of the RBP upregulated in cRIC, 51 (32.5%) of the downregulated in cRIC and 67 (51.1%) of the RBPs detected by vRIC were already linked to virus infection (Figure S7B). Taken together, these results support that our dataset is rich in known regulators of viral infection, but still contains dozens of new host-virus interactions awaiting to be tested experimentally.

**Figure 7.**
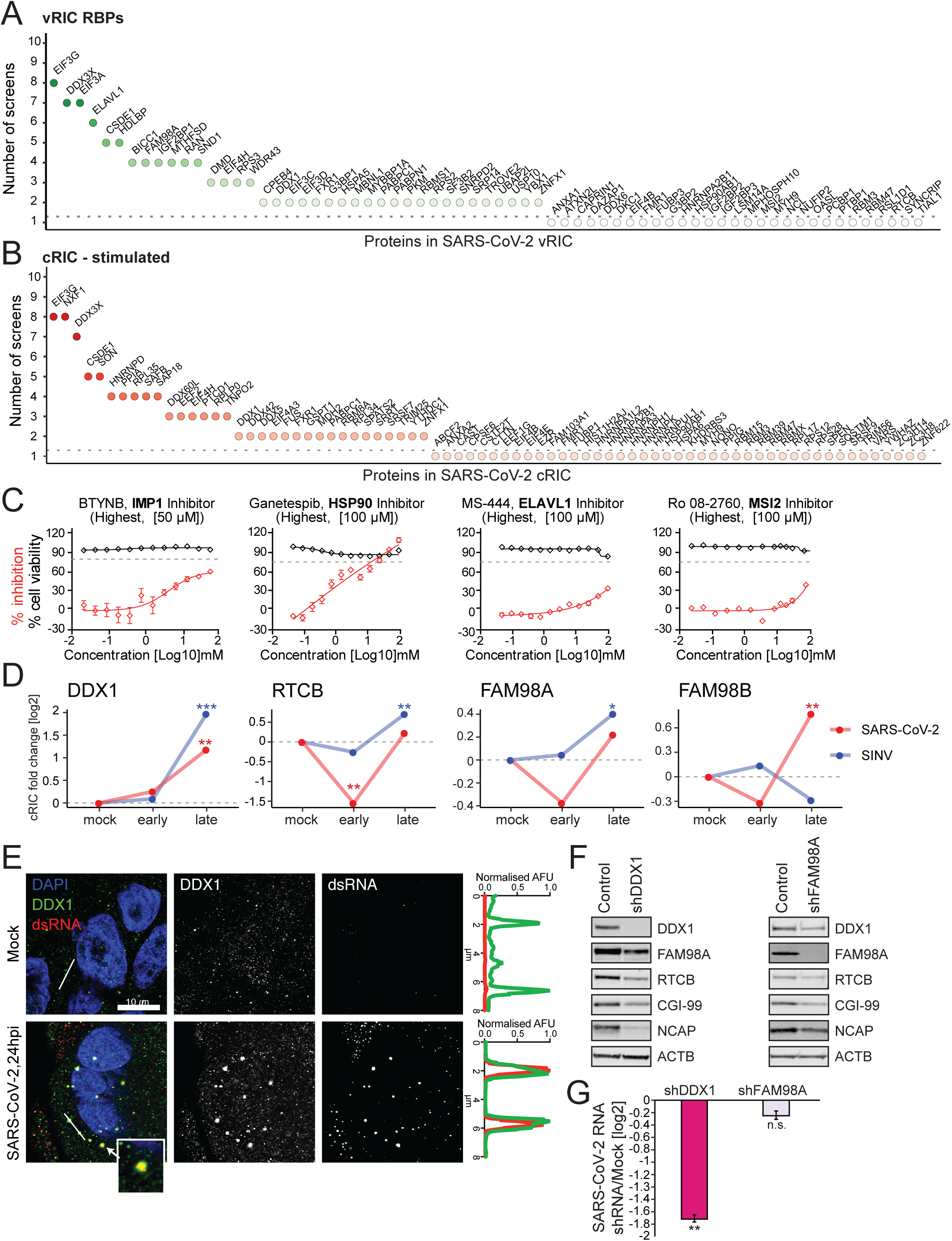
Functional characterisation of protein-RNA interactions in SARS-CoV-2 infected cells. A-B) Proteins with identified phenotypes in genome-wide screens using viruses. RBPs enriched in SARS-CoV-2 vRIC (A) or upregulated in the cRIC experiment (B) are displayed along the x axis, while y axis indicates the number of screens in which the protein has caused a phenotype in infection. C) Effects of RBP inhibitors on SARS-Cov-2 infection. Red line indicates the effects in infection measured by protein ELISA at each drug dose. Black line shows cell viability at each drug dose. Error bars are SEM from three independent experiments. D) RNA binding profiles of the components of the tRNA ligase complex in SARS-CoV-2 (red) and SINV (blue) infected cells (As in Figure 4D) *, FDR < 20%; **, FDR < 10% and *** FDR < 1%. E) Confocal immunofluorescence images of SARS-CoV-2 and mock-infected Calu-3 cells using antibodies against DDX1 and dsRNA. Fluorescence plot shows green and red fluorescence intensity profiles across an 8 µm section (white line). F) Western blot analysis showing the nucleocapsid (NCAP), components of the TLRC complex and β-actin (ACTB) levels in control cells and upon DDX1 or FAM98A knock down. G) SARS-Cov-2 RNA levels in control cells and upon DDX1 or FAM98A knock down measured by RT-qPCR and normalised to β-actin mRNA. Error bars are SEM from three independent experiments.

To determine the biomedical potential of cellular RBPs for COVID-19 treatment, we compared the subset of RBPs stimulated by SARS-CoV-2 infection and the subset of proteins that interact with SARS-CoV-2 RNA to drug databases (Figure S7C). Importantly, 54 proteins within these datasets have potential inhibitors available (Figure S7C). This opens new avenues for the discovery of antivirals targeting RBPs. As a proof of principle, we tested five of these drugs in Calu-3 cells infected with SARS-CoV-2 (Figure 7C and S7D). Our results show that two of these compounds targeting HSP90 and IGF2BP1 (IMP1) cause a strong inhibition of SARS-CoV-2 protein production, with two additional drugs targeting ELAVL1 (HuR) and MSI2 causing moderate effects and one compound targeting PKM having slight effects. Moreover, the inhibitor of PPIA (cyclophilin A) cyclosporine is being tested in the clinic for the treatment of COVID-19 (Rudnicka et al., 2020). These results reflect the potential of protein-RNA interactions as targets for antiviral drugs.

### The tRNA ligase complex, a new regulator of SARS-CoV-2 infection

vRIC revealed DDX1, RTCB and FAM98A as components of the SARS-CoV-2 vRNP. These proteins, together with FAM98B, C14ORF166 (CGI99, CLE) and C2ORF49 (ASW) form the tRNA ligase complex (tRNA-LC) (Popow et al., 2011). DDX1, RTCB, FAM98A and FAM98B interact directly with RNA and are regulated by both SARS-CoV-2 and SINV infection. While DDX1 displayed a continuous increase in RNA-binding activity in the cRIC experiment, the other proteins follow an early inhibition and late increase pattern (Figure 7D). The tRNA-LC mediates the ligation of two unusual RNA fragments, one displaying a 3’-phosphate or 2’, 3’-cyclic phosphate, and the other a 5’-hydroxyl group. This cleavage pattern is generated by a limited repertoire of cellular endonucleases, which include the endoplasmic reticulum resident protein IRE1, which is activated in response to UPR (Jurkin et al., 2014; Popow et al., 2011). Viruses are known to cause UPR, suggesting that they should activate the endonuclease IRE1 (Galluzzi et al., 2017). UPR leads to the tRNA-LC-dependent, cytoplasmic splicing of *Xbp1* mRNA, which encodes a critical transcription factor that coordinates the cellular responses to UPR (Jurkin et al., 2014).

The regulation of the RNA-binding components of the tRNA-LC by SARS-CoV-2 infection and their presence in viral RNPs strongly suggest their involvement in the viral lifecycle. To confirm the interaction between the tRNA-LC and SARS-CoV-2 RNA, we performed an immunofluorescence analysis of infected Calu-3 cells. DDX1, a core component of the tRNA-LC, concentrates at the cytoplasmic foci where dsRNA accumulates, confirming that DDX1 engages with SARS-CoV-2 RNA.

To test the relevance of the tRNA-LC in SARS-CoV-2 infection, we generated A549-ACE2 cells with tetracycline inducible expression of shRNAs against DDX1 and FAM98A. Knocking down of DDX1 led to the depletion of other components of the tRNA-LC, including the ligase RTCB, FAM98A and, in a lesser, extent CIG-99 (Figure 7F and S7E). These results support previous observations showing that the stability of the tRNA-LC relies on presence of the core subunits of the complex (Jurkin et al., 2014). Knock down of the peripheral member of the tRNA-LC, FAM98A, causes minor, non-significant effects in the levels of the other components (Figure S7E). Importantly, silencing of DDX1 caused a strong reduction of intracellular SARS-CoV-2 RNA that correlates with a parallel reduction of NCAP (Figure 7E-F and S7E). Knockdown of FAM98A led to milder effects in both viral RNA levels and NCAP accumulation (Figure 7F-G and S7E). Since DDX1 is a core subunit of the tRNA-LC and FAM98A is a secondary one, these differential effects are expected. Together, these results reveal that tRNA-LC plays an important role in SARS-CoV-2, although its precise mechanism of action requires further investigation.

## OUTLOOK

We provide a systematic and comprehensive analysis of protein-RNA interactions in SARS-CoV-2 infected cells. We show that SARS-CoV-2 infection induces a pervasive remodelling of the RBPome with more than three hundred RBPs displaying enhanced or reduced interaction with RNA. These alterations are not due to changes in protein abundance, but rather due to changes in the cell’s transcriptome and, potentially, posttranslational control. Importantly, nearly 40% of the RBPs that respond to SARS-CoV-2 infection are also regulated by SINV infection, which reflect the existence of prevalent host RBP-virus interactions that span virus species. Similar work with other viruses and cell types will reveal the complement of cellular RBPs with broad participation in virus infection. We also applied a new method to elucidate the composition of SARS-CoV-2 RNPs. Our results reveal dozens of cellular proteins that interact with SARS-CoV-2 RNAs, which are promising for the development of new therapeutic approaches. The relevance and complementarity of our datasets is illustrated by the discovery of the tRNA-LC as key regulator of SARS-CoV-2. Our work thus paves the way to elucidate the importance of the tRNA-LC and other RNA metabolic processes in SARS-CoV-2 biology. Our study also discovers novel viral RNPs, including S, M and ORF9b, opening new angles to investigate their roles in SARS-CoV-2 infection.

In the future, cRIC and vRIC could be extended to other coronaviruses and other biological models such as primary cells and organoids. We are hopeful that this work will further shed light on the pathogenesis of SARS-CoV-2 and accelerate the discovery of therapies for COVID 19.

## Supporting information

cRIC SARS-CoV-2

WCP SARS-CoV-2

SARS-CoV-2 regulated RBPome

SARS-CoV-2 RBP response profiles

vRIC SARS-CoV-2

WCP cRIC vRIC Normalisation

Genome wide screens summary cRIC vRIC

annotation-definitions

## ACKNOWLEDGEMENTS

We thank our friend Bernd Fischer, who sadly passed away, for his contribution to the develop RBDetect. We also thank Marie Bartenschlager for excellent technical support in SARS-CoV-2 related assays. We thank Catarina Franco for her assistance in the operation of the Orbitrap Eclipse mass spectrometer. Alfredo Castello is supported by MRC Career Development Award MR/L019434/1, MRC grants MR/R021562/1, MC_UU_12014/10 and John Fell funds from the University of Oxford. Ralf Bartenschlager is supported by the Deutsche Forschungsgemeinschaft (DFG, German Research Foundation) – Project number 240245660 – SFB 1129, project number 272983813 – TRR 179 and the German Center for Infection Research (DZIF), project numbers 8029801806 and 462 8029705705. Ilan Davis is funded by the Wellcome Investigator Award 209412/Z/17/Z and Wellcome Strategic Awards 091911/B/10/Z and 107457/Z/15/Z. Javier Martinez is funded by the Medical University of Vienna, the “Fonds zur Förderung der wissenschaftlichen Forschung (FWF)” as Stand-Alone Project P29888, and through the RNA Biology Doctoral Program, FWF. JYL is funded by the Medial Sciences Graduate Studentship, University of Oxford. Wael Kamel is funded by the European Union’s Horizon 2020 research and innovation programme under Marie-Sklodowska-Curie grant agreement 842067. Anna Andrejeva is supported by European Union Horizon 2020 INFRAIA project Epic-XS (project no. 823839). Michael Knight is funded by the BBSRC DTP scholarship number BB/M011224/1. We thank Olympus UK and Europe for their generous loan of an Olympus ScanR-Sora spinning disc microscope for the imaging carried out for this project.

## TABLE LEGENDS

**Table S1: Analysis of RBP dynamics in SARS-CoV-2 infected cells by cRIC.** cRIC datasets were generated at 8 and 24 hpi.

**Table S2: Analysis of the total proteome of SARS-CoV-2 infected or mock-infected cells.** WCP analyses were generated at 8 and 24 hpi.

**Table S3: Post-translational modification mapping to cellular RBPs.** The cRIC data was cross-referenced to phospho-proteomic and ubiquinomic datasets reported by Bouhaddou et al., 2020; Klann et al., 2020; Stukalov et al., 2020.

**Table S4: Classification of RBPs by RNA-binding responses to SARS-CoV-2 infection.** The different protein clusters were obtained by comparing the 8hpi/mock and 24hpi/8hpi fold-change of each RBP.

**Table S5. vRIC analysis of the SARS-Cov-2 RNA interactome.**

**Table S6. cRIC and vRIC normalisation to the WCP.** This table provides a ‘cross-linkability’ index that can be used to classify RBPs based on their ability to crosslink to RNA.

**Table S7. RBP potential for regulatory roles in virus infected cells.** List of RBPs reported in this study linked to the genome-wide screens that have reported their involvement in virus infection.

**Table S8. Antibodies and Oligonucleotide primers used in this study.**

**Table S9. List of GO-terms and KEGG pathways related to immune response and known RNA binding domains.**

## MATERIALS AND METHODS

### Experimental model and subject detail

#### Cell culture

Calu-3 cells (kind gift from Dr. Manfred Frey, Mannheim, Germany) were maintained in DMEM (Gibco, 41965039) with 20% fetal bovine serum (FBS) (Gibco, 10500064) and 1x penicillin/streptomycin (Sigma Aldrich, P4458) at 37°C with 5% CO_2_. A549-Ace2 (Klein et al., 2020) were maintained as above with 10% FBS. To generate inducible knockdown lines, cells were infected with Lentiviral vectors derived from pLKO-Tet-On (Wiederschain et al., 2009) with the guide sequence GATGTGGTCTGAAGCTATTAA for DDX1 and GCACATTCAGTAGCCTTATTT for FAM98A. Lentiviruses were produced by co-transfection of HEK293T cells with pHEF-VSVG (NIH AIDS Research & Reference reagent program #4693) and psPAX2 (kind gift N. Proudfoot, Oxford, UK). After infection of A549-Ace2 cells with the lentiviruses, selection was performed with 1 µg/ml. shRNAs were induced by addition of 1ug/ml doxycycline.

#### Viruses

Infection of Calu-3 cells for virus growth kinetics, cRIC, vRIC, WCP and drug screen was performed using isolate hCoV-19/Germany/BavPat1/2020 (European Virology Archives: 026V-03883, EPI_ISL_406862). For validation in knockdown studies and immunofluorescence, hCoV-19/England/02/2020 (Public Health England propagated viral isolate Feb 2020, EPI_ISL_407073) was used.

### Virus growth kinetic experiments

1.2 x 10^5^ Calu-3 cells were seeded into each well of a 24-well plate. Cells were infected 24 hours after seeding with SARS-CoV-2 at a multiplicity of infection (MOI) of 1. To determine infectivity, 50 µl of supernatant from each well was used in plaque assays. Plaque assays were performed as previously described (Klein et al., 2020). Briefly, 2.5 x 10^5^ Vero cells were seeded into each well of a 24-well plate and cells were inoculated with 10-fold serial dilutions of SARS-CoV-2 containing supernatants for 1 h at 37°C. After 1h, viral supernatants were replaced by serum-free MEM (Gibco #11095080, Life Technologies) containing 0.8% carboxymethylcellulose (Sigma, 11095080). Three days later, plates were fixed with 6 % formaldehyde for 30 minutes and rinsed with tap water. Plates were stained with a solution containing 1% crystal violet (Sigma, HT90132-1L) and 10% ethanol for 30 min. After rinsing with tap water, plaques were counted to determine viral titer.

For intra- and extra-cellular RNA extraction, NucleoSpin RNA extraction kit (Macherey-Nagel, #740955.50) was used following the manufacturer’s specifications. cDNA synthesis from the total RNA isolated was achieved using a high-capacity reverse transcription kit (ThermoFisher, #4368814). cDNA samples were diluted 1:15 and used for qPCR with the iTaq Universal SYBR green mastermix (Biorad, #1725120). Cycle threshold values were corrected for PCR efficiency of each primer set and normalized to the hypoxanthine phosphoribosyltransferase 1 (HPRT) mRNA to determine relative abundance of viral RNA for each sample (see table S8).

### Cell viability assay and determination of infection rate

To establish cell viability and infection rate, 1.2 x 10^5^ Calu-3 cells were seeded into each well of a 24-well plate onto glass coverslips. Mock-infected and SARS-CoV-2-infected cells were fixed at the times post infection indicated in the figures with 6% formaldehyde for 30 min. Cells were washed twice with PBS (phosphate-buffered saline) and permeabilized with 0.2% Triton X-100 in PBS. Permeabilized samples were incubated with blocking solution (2% of milk and 0.02% Tween-20 in PBS) for 1 h at room temperature. Samples were stained with primary antibodies specific to dsRNA (see Table S8) as well as DAPI (DAPI Fluoromount-G, SouthernBiotech, 0100-20) to visualize the nuclei using a Nikon Eclipse Ti microscope (Nikon, Tokio, Japan). Three replicates per time point were analyzed. Nuclei were counted with a custom-made macro for the Fiji software package (Schindelin et al., 2012). Number of nuclei in infected samples were normalized to the non-infected control counterparts. To determine the infection rate, the number of infected cells at each time point was determined using the dsRNA fluorescence signal with Fiji software using a custom macro (Schindelin et al., 2012).

### Colorimetric cell-based assay to assess the effects of RBP inhibitors in SARS-CoV-2 infection

Calu-3 cells were seeded at 2 x 10^4^ cells per well of 96-well plate. Cells were treated 24 hours later with 2-fold serial dilutions of the indicated compounds in duplicate wells. Dilutions ranged from 2.5 nM to 50 µM for Ro 08-2750 (TOCRIS, #2272) and the BTYNB IMP1 inhibitor (Cayman Chemical, #25623), 5 nM to 100 µM for Ganetespib (BIOZOL, BYT-ORB181166) and MS-444 (Hycultec, HY100685-1mg) and 1,25 nM to 25 µM for the PKM2 inhibitor - compound 3k (BIOZOL, SEL-S8616)). 2 hours after treatment, cells were infected with SARS-CoV-2 (BavPat1/2020 strain) at a MOI of 2. At 24 h post infection, plates were fixed with 6% formaldehyde for 30 min. Cells were then washed twice with PBS (Phosphate-buffered Saline) and permeabilized with 0.2% Triton X-100 in PBS. Permeabilized samples were then incubated with blocking solution (2% of milk and 0.02% Tween-20 in PBS) for 1 h at room temperature. Blocking solution was replaced with primary antibodies specific for SARS-CoV NCAP (Table S8) diluted in blocking solution. Cells were incubated for 1 h at 37 °C, washed four times with PBS followed by incubation with horse radish peroxidase (HRP)-conjugated secondary antibodies diluted in PBS (containing 0.02% Tween-20) for 1 h at 37 °C. Wells were washed 3 times with PBS. PBS excess was carefully removed, and wells were developed by adding 50 µl of TMB Microwell Peroxidase (SeraCare, Cat: 5120-0077) to each well for 5 min followed by 50 µl of 0.5 M of H_2_SO_4_ solution to stop the reaction. Absorbance was measured at 450 nm using a Tecan-Sunrise absorbance microplate reader. Values were normalized to vehicle (DMSO). In order to assess the effects of the above-mentioned inhibitors on cell viability, we employed the commercial kit CellTiterGlo® Luminescent Cell Viability Assay (Promega, Cat: G7570) on a Mithras LB 940 plate reader (Berthold Technologies). The assays were performed following the manufacturer’s instructions in uninfected cells for the different doses of each compounds. Luminiscence values were normalised to vehicle (DMSO).

### Immunofluorescence

Round #1.5 (diameter 13 mm) coverslips (Thermo Fischer Scientific) were wiped with lint-free tissue soaked in 80% ethanol and washed in 100% ethanol twice for 2 h. 2×10^5^ Calu-3 cells were seeded on the dried coverslips and incubated in growth media for 48 hours prior to the experiment. Cells were infected with 2×10^5^ PFU/well (MOI=1) SARS-CoV-2 (hCoV-19/England/02/2020) and incubated for 24 hours. Cells were fixed in 4% formaldehyde for 30 minutes and washed once with PBS. Cells were permeabilised for 10 min with PBSTx (1x PBS + 0.1% Triton X-100) at room temperature. Next, cells were washed twice in PBSTw (1x PBS + 0.1% Tween-20) for 5 min each and incubated in blocking solution (PBSTw + 2.5% goat serum + 2.5% donkey serum) for 1 h at room temperature. Cells were incubated overnight at 4°C with primary antibodies diluted in blocking solution (table S8). Coverslips were then washed three times with PBSTw for 10 min each at room temperature and incubated with secondary antibodies and DAPI (1 µg/ml) diluted in blocking solution overnight at 4°C. Cells were washed three times with PBSTw for 10 min each, once in PBS for 10 min, once in milliQ H_2_O and the coverslips were mounted on glass slides using Vectashield HardSet mounting medium (Vector Laboratories #H-1400). Mounted cells were imaged on an Olympus SoRa spinning disc confocal with Orca Flash4 CMOS camera using 100x silicone oil objective (1.35 NA, UPLSAPO100XS). Specimens were imaged in at least six different locations per coverslip. 3D-stacked images were taken with voxel size of 80 nm x 80 nm x 200 nm in x:y:z and images were deconvolved with maximum likelihood algorithm using cellSens (5 iterations, default PSF, no noise reduction, Olympus). Background subtraction was performed on all channels using rolling ball subtraction method (radius = 250 px) in ImageJ (National Institutes of Health). Fluorescence intensity profiles were obtained using ImageJ “Plot profile” tool across 8 µm regions on 0.4 µm max intensity z-projected images. Voxel intensities were normalized to maximum intensity value obtained from ‘SARS-CoV-2 infected’ condition.

### Comparative RNA interactome capture

Comparative RNA interactome capture (cRIC) was performed based on the previously described protocol (Castello et al., 2013) with the following alterations: Calu-3 cells were grown in sets of 3×15 cm dishes with 10^7^ cells/dish. One set of dishes remained uninfected while a second set was infected with SARS-CoV2 (hCoV-19/Germany/BavPat1/2020) at a MOI of 1. One of these infected cell sets was incubated for 8 h and the other for 24 h. 3 biological replicates for each condition were performed. After incubation, plates without lids were placed on ice and cells were irradiated with 150 mJ/cm^2^ of UV light at 254 nm and lysed with 5 mL of lysis buffer (20 mM Tris-HCl pH 7.5, 500 mM LiCl, 0.5% LiDS wt/vol, 1 mM EDTA, 0.1% IGEPAL (NP-40) and 5 mM DTT). Lysates were homogenized by passing the lysate at high speed through a 5 mL syringe with a 27G needle, repeating this process until the lysate was fully homogeneous. Ten percent of the lysate was separated for total proteome analysis (WCP). The rest of the samples were processed as follows. Protein content was measured using Qubit protein assay (Invitrogen Q33212) and lysates were normalized by protein content. 0.45 mL of pre-equilibrated oligo(dT)25 magnetic beads (New England Biolabs, #S1419S) were added to the lysates and incubated for 1 h at 4°C with gentle rotation. Beads were collected in the magnet and the lysate was transferred to a new tube and stored at 4°C. Beads were washed once with 5 mL of lysis buffer, followed by two washes with 5 mL of buffer 1 (20 mM Tris-HCl pH 7.5, 500 mM LiCl, 0.1% LiDS wt/vol, 1 mM EDTA, 0.1% IGEPAL and 5 mM DTT), and two washes with buffer 2 (20 mM Tris-HCl pH 7.5, 500 mM LiCl, 1 mM EDTA, 0.01% IGEPAL and 5 mM DTT), in all cases incubating the beads for 5 min at 4°C with gentle rotation.

Beads were then washed twice with 5 mL of buffer 3 (20 mM Tris-HCl pH 7.5, 200 mM LiCl, 1 mM EDTA and 5 mM DTT) at room temperature for 3 minutes. Beads were resuspended in 300 μL of elution buffer and incubated for 3 min at 55°C with agitation. After collecting the beads with a magnet, eluates (supernatants) were collected and stored at −80°C. The lysates were subjected to a second round of capture and the eluates from the first and second cycles were combined. Prior to mass spectrometry sample processing, samples were RNase treated with ∼0.02U RNase A and RNase T1 at 37°C for 1h.

### Viral RNA interactome capture

Viral RNA interactome capture was performed as in (Kamel et al., In preparation). Briefly, Calu-3 cells were grown in sets of 2×15 cm dishes. For the infected samples (SARS-CoV2/4SU/Fvo), at 8hpi (hours post-infection, MOI=1), the growth media were replaced with fresh media supplemented with (20 µM Flavopiridol hydrochloride hydrate (Fvo, Cat.No. F3055, Sigma-Aldrich)) and 100 µM 4-Thiouridine (4SU, Cat.No. T4509, Sigma-Aldrich)). The plates were returned to the incubator for additional 16 hours. At 24hpi, growth media were discarded, and the cells were rinsed once with PBS (Phosphate-buffered saline). Cells were irradiated twice with at 200 mJ/cm^2^ using ultraviolet light 365nm. At this stage, samples were subjected to the standard RNA-interactome capture described above. For the control uninfected samples (M/4SU/Fvo), cells were treated as in SARS-CoV-2/4SU/Fvo with exception of not adding the virus. Both M/4SU/Fvo and SARS-CoV-2/4SU/Fvo were performed in sets of four biological replicates. Additional controls, (M/4SU/-), uninfected cells were treated as in M/4SU/Fvo, without the addition of Fvo, and (M/-/-) uninfected cells were incubated with growth media (not supplemented with Fvo and 4SU) and not crosslinked. Both (M/4SU/-) and (M/-/-) were performed in sets of three biological replicates.

### Mass spectrometry

The cRIC, vRIC and WCP protein samples were processed via the bead-based single-pot, solid-phase-enhanced sample-preparation (SP3) method, using Speed Bead Magnetic Carboxylate Modified Particles (Sigma-Aldrich, cat.no.45152105050250) (Hughes et al., 2019). Protein digestion was performed using Trypsin Gold (MS grade; Promega, cat. no. V5280). Processed peptides were acidified by formic acid (final concentration 5%) prior to Mass spectrometry analysis.

For cRIC and vRIC peptides, liquid chromatography (LC) was preformed using Ultimate 3000 ultra-HPLC system (Thermo Fisher Scientific). Peptides were initially trapped in C18 PepMap100 pre-column (300 µm inner diameter x 5 mm, 100A, Thermo Fisher Scientific) in Solvent A (Formic acid 0.1% (v/v), Medronic acid 5 µM). Trapped Peptides were separated on the analytical column (75 µm inner diameter x 50cm packed with ReproSil-Pur 120 C18-AQ, 1.9 mm, 120 A, Dr. Maisch GmbH) in a 60min 15%–35% [vol/vol] acetonitrile gradient with constant 200 nL/min flow rate. Eluted peptides were directly electrosprayed into a QExactive mass spectrometer (Thermo Fisher Scientific). Mass spectra were acquired in the Orbitrap (scan range 350-1500 m/z, resolution 70000, AGC target 3 x 10^6^, maximum injection time 50 ms) in a data-dependent mode. the top 10 most abundant peaks were fragmented using CID (resolution 17500, AGC target 5×10^4^, maximum injection time 120 ms) with first fixed mass at 180 m/z.

Both WCP and vRIC peptides were analysed using a Dionex Ultimate 3000 RSLC nanoUPLC (Thermo Fisher Scientific Inc, Waltham, MA, USA) system online with an Orbitrap Eclipse mass spectrometer (Thermo Fisher Scientific Inc, Waltham, MA, USA). Peptides were loaded onto a trap-column (Thermo Scientific PepMap 100 C18, 5 μm particle size, 100A pore size, 300 μm i.d. x 5mm length) and separation of peptides was performed by C18 reverse-phase chromatography at a flow rate of 300 nL/min and a reverse-phase nano Easy-Spray column (Thermo Scientific PepMap C18, 2μm particle size, 100A pore size, 75μm i.d. x 50cm). WCP peptides were acquired in a 120 min run while vRIC samples in an 82 min run. Analytical chromatography for WCP peptides consisted of Buffer A (0.1% formic acid in HPLC-grade water) and Buffer B (80% ACN, 0.1% formic acid). 0-3 min at 2% buffer B, 3-90 min linear gradient 2% to 40% buffer B, 90-90.3 min linear gradient 40% to 90% buffer B, 90.3-95 min at 90% buffer B, 95-95.3 min linear gradient 90% to 2% buffer B and 95.3-120 min at 2% buffer B. Analytical chromatography for vRIC peptides was Buffer A (HPLC H_2_O, 0.1% formic acid) and Buffer B (80% ACN, 0.1% formic acid). 0-3 min at 3.8% buffer B, 3-63 min non-linear gradient 3.8% to 40% buffer B, 63-63.3 min linear gradient 40% to 90% buffer B, 63.3-68 min at 90% buffer B, 68-68.3 min non-linear gradient 90% to 3.8% buffer B and 68.3-82 min at 3.8% buffer B. All *m/z* values of eluting peptide ions were measured in an Orbitrap mass analyzer, set at a resolution of 120 000 and were scanned between *m/z* 380-1500 Da. Data dependent MS/MS scans (3 second duty cycle time) were employed to automatically isolate and fragment precursor ions using Collisional-induced Dissociation (CID) (Normalised Collision Energy of 35%). Only precursors with charge between 2 to 7 were selected for fragmentation, with an AGC target and maximum accumulation time of 1×10^4^ and 125 ms respectively. Precursor isolation was performed by the quadrupole with 1.2 m/z transmission window. MS2 fragments were measured with the Ion Trap analyser. Dynamic exclusion window was set to 70 seconds.

Protein identification and quantification were performed using Andromeda search engine implemented in MaxQuant (1.6.3.4) (Cox 2011) under default parameters (Cox et al., 2011). Peptides were searched against reference Uniport datasets: human proteome (Uniprot_id: UP000005640, downloaded Nov2016) and SARS-CoV-2 (Uniprot_id: UP000464024, downloaded 24June2020). False discovery rate (FDR) was set at 1% for both peptide and protein identification. For cRIC and WCP samples, MaxQuant search was performed with “match between run” activated. For vRIC samples, since each sample was analyzed on both Eclipse and QExactive mass spectrometers, raw spectra form both runs were combined as separate fractions in the MaxQuant search (the spectra from the Eclipse was assigned fraction 1 and the spectra from the QExactive is assigned fraction 5, and each sample was as independent experiment).

### Proteomic quantitative analysis

For relative quantification, MaxQuant outputs (proteinGroups) were used for downstream analysis. Proteins flagged as potential contaminants were filtered out, using R-package “DEP (1.4.1)”, together with proteins with all missing values. In case of cRIC and WCP experiments, proteins raw intensities were normalized and transformed using R-package Variance Stabilizing Normalization “VSN (3.50.0)”. Correlation analysis between replicates was preformed using R-package “PerformanceAnalytics (v2.0.4)”. Missing value imputation was only preformed for proteins with missing values in all replicates in one experimental condition, while present in the other condition (at least in 2 out of 3 replicates). Imputation was preformed using local (by sample) minimum determination method (Mindet) (Lazar et al., 2016). Statistical analysis for the processed intensities was performed in R-package “limma (3.38.3)” using empirical Bayesian method moderated t-test. P values were adjusted for multiple-testing using Benjamini-Hochberg method. For the vRIC experiments, samples were processed as described above with exception of the normalization step.

### Clustering of cRIC responses to SARS-CoV-2 infection

Cellular RBP responses to SARS-CoV-2 infection was classified into initial response, which is defined as cRIC log fold change from mock to early time point post-infection (8 hpi/mock), and progressive response, which is determined by log fold change from early to late time point post-infection (8 hpi/24 hpi). Protein abundance fold changes in these two stages were visualised using a scatter plot. The RNA-binding activities of cellular RBPs were divided into 8 clusters based on their initial response, progressive response, and FDR. Clustering was based on an FDR<10% with a log(2) fold change of 0.5 as thresholds. For clustering of spliceosome/ spliceosome–related proteins, list of different classes of spliceosomal proteins obtained from Spliceosomedb.

### SARS-CoV-2 proteins RNA binding prediction

RNA binding prediction for regions on the viral protein sequence was performed with RBDetect. RBDetect is a machine learning model trained by Shrinkage Discriminant Analysis (SDA) with a dataset of 8891 experimentally identified polypeptides from the RBDmap experiment, using positive examples (RNA-bound polypeptides) and negative examples (RNA released polypeptides) (Castello et al., 2016). For each amino acid position on the viral protein sequence, RBDetect assigns a probability value to bind RNA based on the fragment centered at that position. Then, a Hidden Markov model is used to visualize the probabilities in a sequential manner, which helps to determine the most probable binding regions on a larger scale.

### Analysis of the PTM profile of RBPs identified in cRIC

Cellular RBPs detected in cRIC experiments were cross-referenced to phosphorylation and ubiquitination sites of recent large scale (post-translation modification) PTM quantification experiments preformed in SARS-CoV-2 infected cells. PTM datasets obtained from the single-timepoint phospho-proteome work by (Klann et al., 2020), multi-timepoints phospho-proteome work by (Bouhaddou et al., 2020), and multi-level omics work by (Stukalov et al., 2020). The SARS-CoV-2 regulated RBPome is defined as RBPs with FDR < 0.1 in the cRIC experiment. SARS-CoV-2 regulated PTM sites are significant hits in each data sources using the criteria defined in the corresponding publications. For the multi-timepoint dataset, a PTM site is considered SARS-CoV-2 regulated, if it is determined as significant at any timepoint. Fisher’s exact test was employed to calculate odds ratios and significance of enrichment of each PTM annotation in the SARS-CoV-2 upregulated RBPome versus downregulated RBPome.

### Drug-protein interactions

Cellular RBPs (stimulated upon SARS-CoV-2 infection (from cRIC) or bound to viral RNA (from vRIC)) were examined for known chemical compound interactions through the Drug-Gene Interaction database (DGIdb, downloaded Oct-2020).

### Gene Ontology (GO) terms

Using the GO annotation available via the GO.db R package (3.11.4), GO terms including the term ‘RNA binding’ (to annotate RNA-binding related functions, processes, or compartments) or term ‘immun’ or exact terms ‘immune response’ and ‘innate immune response’ (to annotate immunity related functions, processes, or compartments) were selected. The full list of terms is provided as a supplementary table (Table S9). The R package org.Hs.eg.db (3.11.4) was used to identify the genes (proteins) in our dataset that are annotated to these GO terms using the cross-database id mapping functionality. GO enrichment analysis was performed using PANTHER classification system (http://www.pantherdb.org).

### Kyoto Encyclopedia of Genes and Genomes (KEGG) pathways

KEGG pathways under the ‘Immune system’ category in the high-level KEGG hierarchy available via the R package “KEGGREST” (1.28.0) were selected (see tableS9) and genes mapping to these pathways were identified using “org.Hs.eg.db.”

### Pfam RNA-binding domains

Classification of proteins into classical and non-classical RNA-binding proteins is based on their Pfam domain composition. We considered RRM, KH, DSRM, Piwi, DEAD, PUF, CSD, and zf-CCCH domains as classical. These were obtained from the PFAM.db R package (3.11.4). Furthermore, we considered as non-classical RNA-binding domains those Pfam-A domains robustly identified as RNA-binding by RBDmap with at least 3 peptides and RNA interactome capture (Castello et al., 2012; Castello et al., 2016). The classification is provided in Table S9. The proteins containing these domains were identified using org.Hs.eg.db.

### PubMed literature linking genes to viral infections

To automatically query the NCBI Entrez Utilities REST API, the R package “rentrez” (1.2.2) was used. For each gene symbol in our dataset the number of PubMed articles matching with a search query “(SYMBOL) AND (virus)” where SYMBOL is the gene name, such as EIF4E were retrieved. A minimum of five search results was considered a substantiated indication of a gene having a connection to virus-related literature.

### RNA sequencing analysis

Strand-specific, poly(A) RNA-seq corresponding to SARS2-infected (MOI=2) Calu-3 cells and controls from published work (Blanco-Melo et al., 2020) were downloaded from the Sequence Read Archive using “SRA toolkit “(2.10.8). Specifically, we analysed the following samples: Calu3 Mock 1 (GEO GSM4462348, SRA series SRX8089276, SRA run SRR11517744), Calu3 Mock 2 (GEO GSM4462349, SRA series SRX8089277, SRA run SRR11517745), Calu3 Mock 3 (GEO GSM4462350, SRA series SRX8089278, SRA run SRR11517746), Calu3 SARS-CoV-2 1 (GEO GSM4462351, SRA series SRX8089279, SRA run SRR11517747), Calu3 SARS-CoV-2 2 (GEO GSM4462352, SRA series SRX8089280, SRA run SRR11517748), Calu3 SARS-CoV-2 3 (GEO GSM4462353, SRA series SRX8089281, SRA run SRR11517749). Raw reads alignment was performed via “STAR aligner” (2.7.3a), with splicing-aware settings, against human reference genome (GRCh38.99) and SARS-CoV-2 (NC_045512.2). Only uniquely aligned reads were used for downstream analyses. Mapped reads (exonic regions) counting was performed by “htseq*-*count” (0.11.3) in a strand-specific fashion. In order to assess the main driver(s) of variations across the RNA-seq samples, we performed a principal component analysis (PCA). First, we performed library size correction and variance stabilisation with regularized–logarithm transformation implemented in “DESeq2” (1.28.1). This corrects for the fact that in RNA-seq data, variance grows with the mean and therefore, without suitable correction, only the most highly expressed genes drive the clustering. The 500 genes showing the highest variance were used to perform PCA using the “prcomp” function implemented in the base R package “*stats”* (4.0.2). Finally, differential expression analysis was performed using the R package “DESeq2” (1.28.1). “DESeq2” estimates variance-mean dependence in count data from high-throughput sequencing data and tests for differential expression based on a model using the negative binomial distribution.

### Data wrangling and visualisation

The “tidyverse suite” (1.3.0) was used for data wrangling in R, and “rtracklayer” (1.48.0) for manipulating gtf annotation files. Furthermore, we used the following R packages in creating the presented visualisation: “ggplot2*”* (3.3.2), “viridis*”* (0.5.1), “ggrepel*”* (0.8.2), “scales” (1.1.1).

**Table 1.**
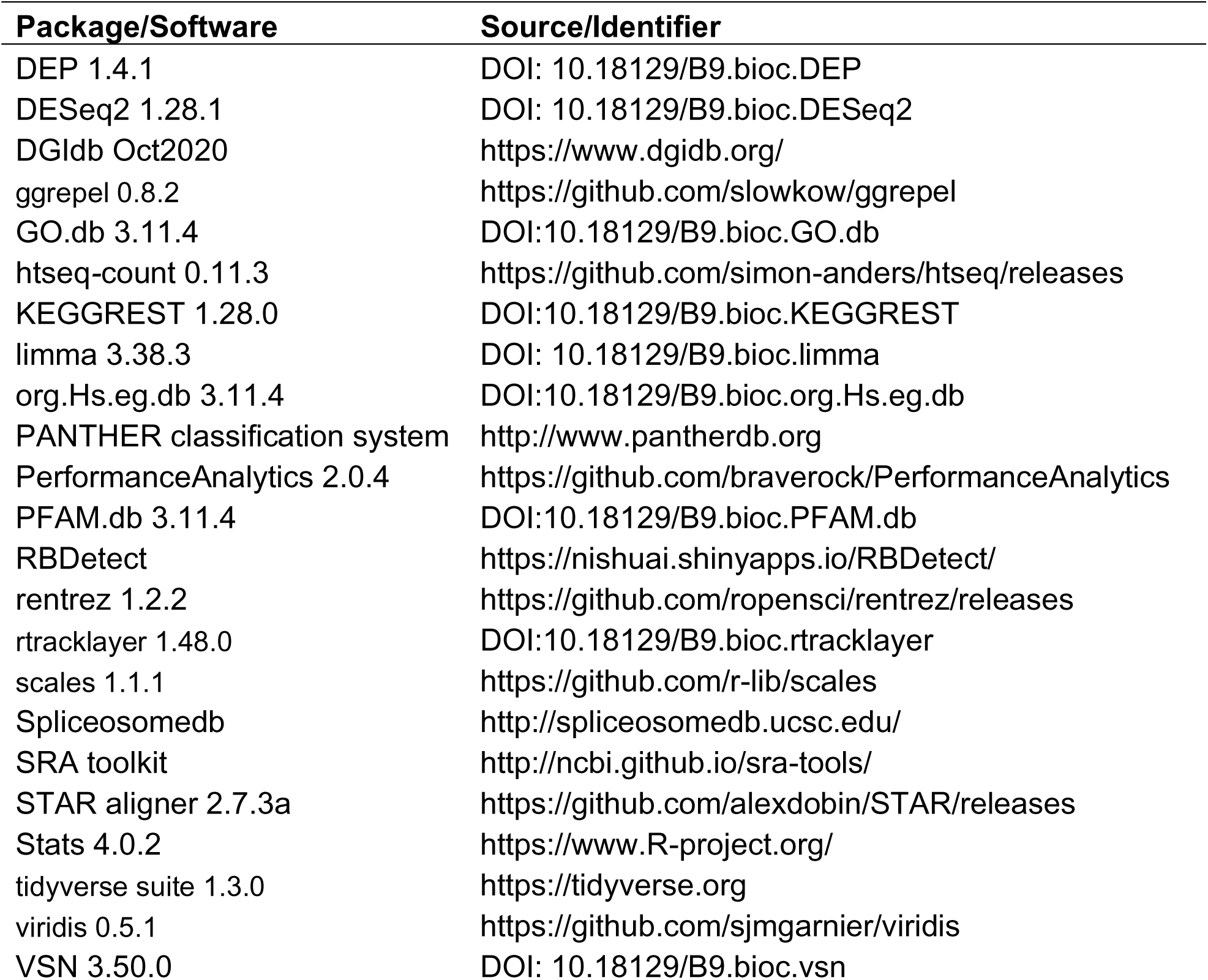

---

**Figure S1:**
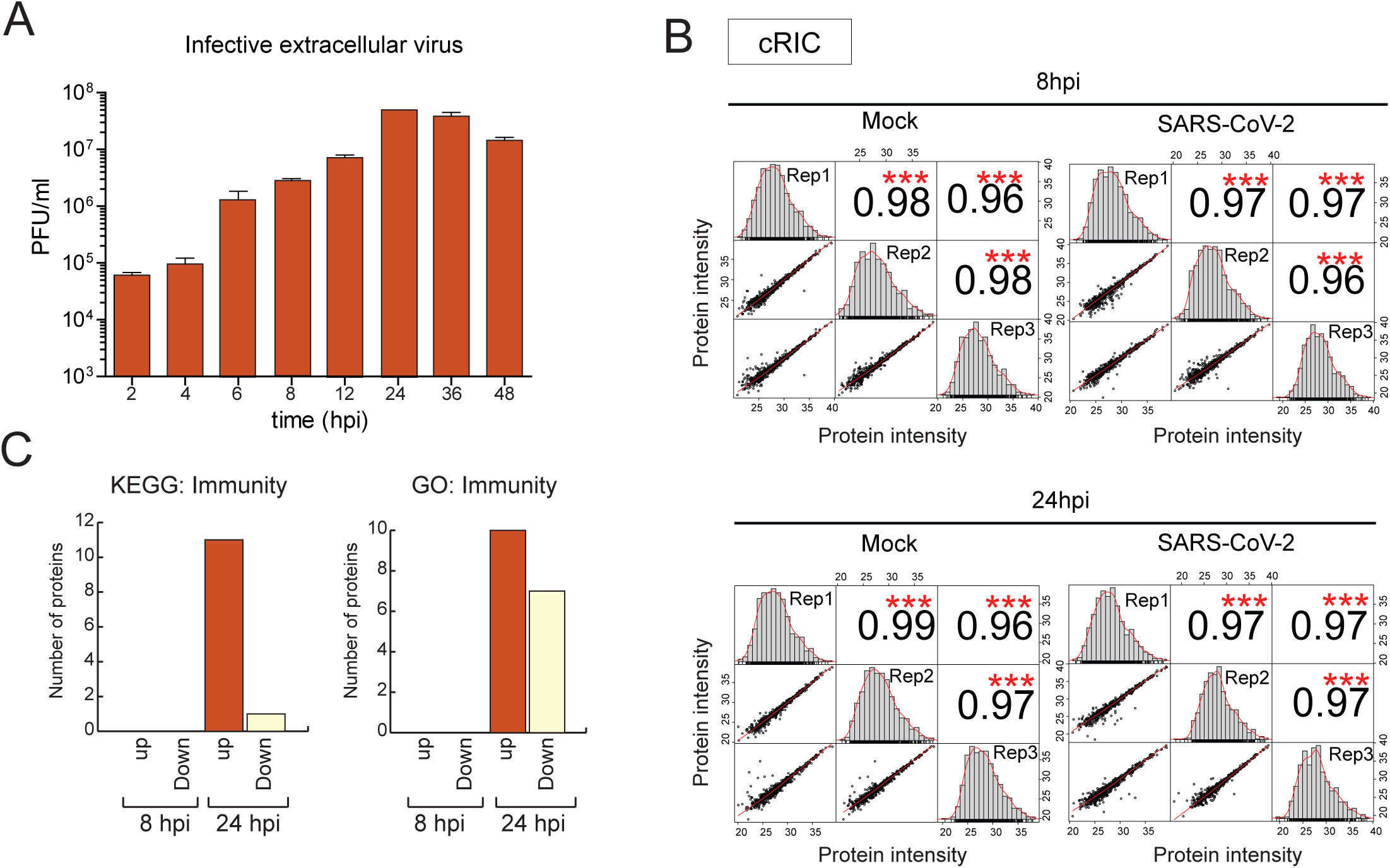
Profiling RBP dynamics by comparative RIC. A) Supernatant of cells infected with SARS-CoV-2 for different times, were collected and titrated by plaque assay. B) Scatter plots comparing protein intensity [log2] across replicates of the total proteome analysis and the different conditions. Pearson correlation is indicated. ***, p<0.001. C) Number of upregulated or downregulated RBPs with annotation related to immunity in KEGG (left) or gene ontology (GO, right).

**Figure S2:**
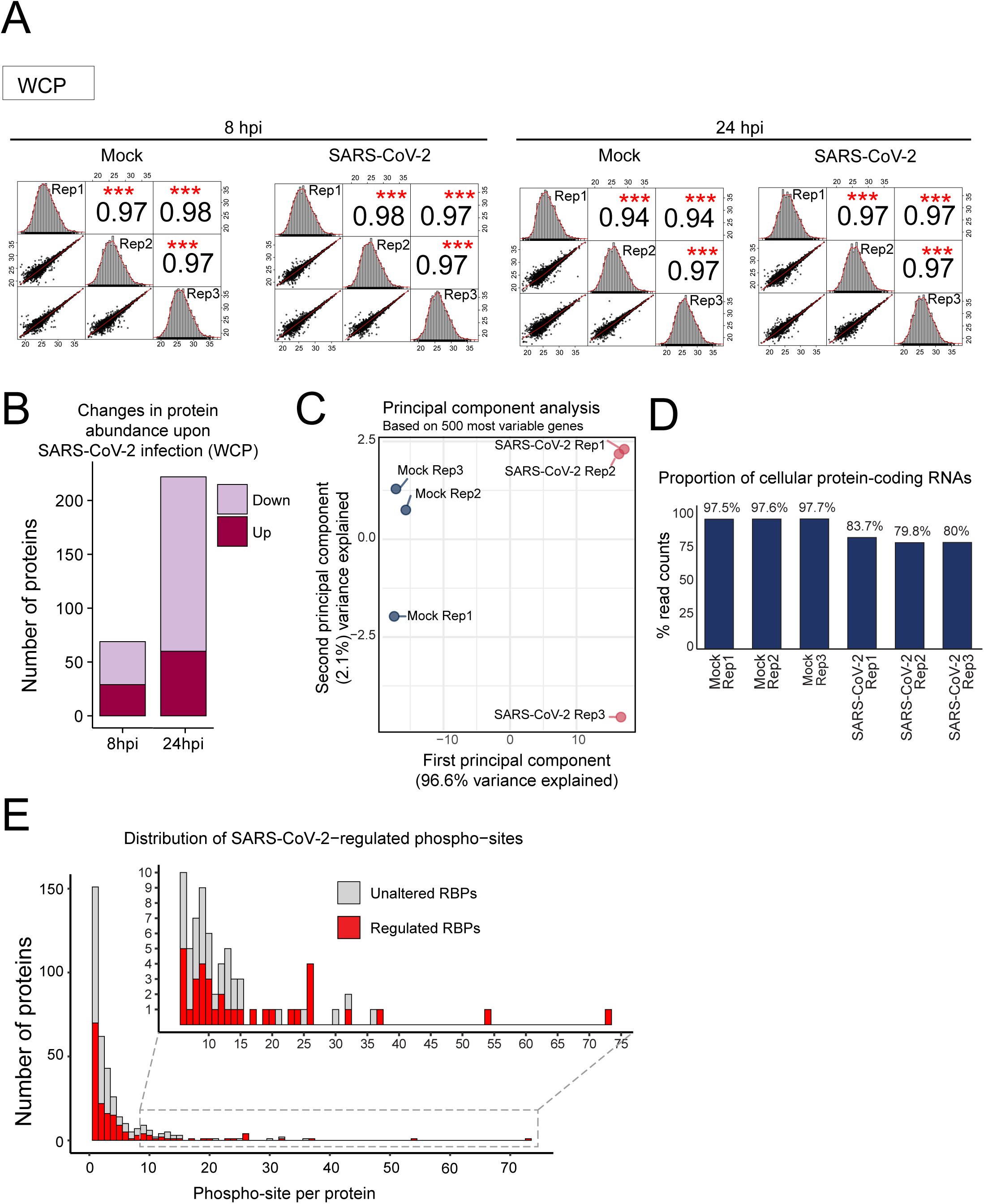
SARS-CoV-2 induced alterations in the whole cell proteome. A) Scatter plots comparing protein intensity [log2] across different replicates. Pearson correlation is indicated. ***, p<0.001. B) Bar-plot showing the proportion of proteins with changes in protein levels upon 8h or 24h of SARS-Cov-2 infection. C) First two components of a principal component analysis (PCA) performed on the 500 genes showing the highest variance in RNA-seq. The first component clearly separates infected cells from uninfected cells (mock in Blue and infected cells in Red). D) Percent of RNA-seq reads assigned to human protein coding genes of total count of uniquely assigned reads. E) Distribution of the number of phosphosites detected in regulated and unaltered RBPs.

**Figure S3.**
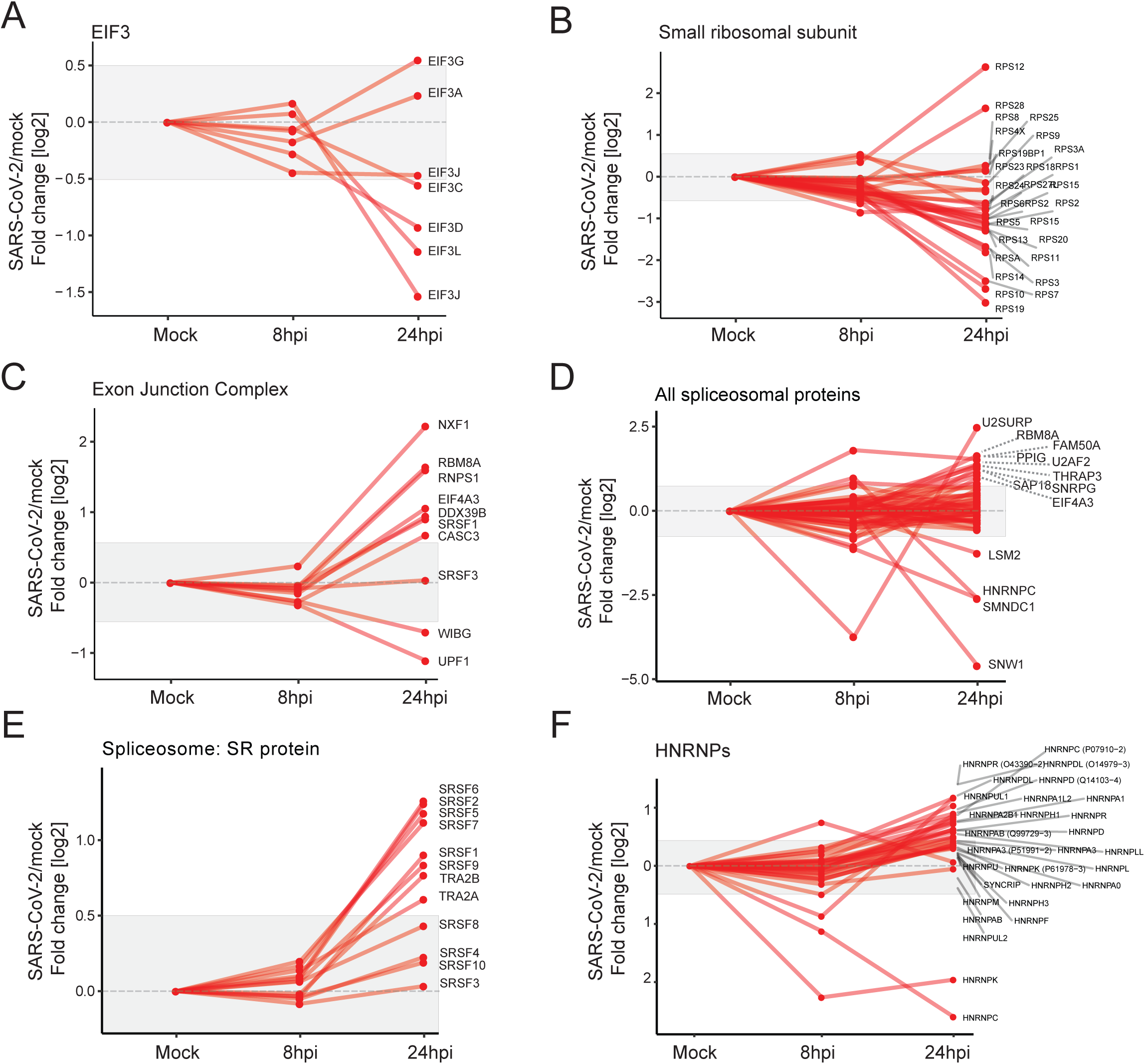
RNA-binding dynamics of functionally related RBPs in response to SARS-coV-2 infection. A-F) Line plots showing the protein intensity ratio between 8hpi/mock and 24hpi/mock samples from the cRIC experiment for functionally related proteins, including EIF3 complex (A), small ribosomal subunit (B), exon junction complex (C), spliceosome (D), SR proteins (E) and HNRNPs (F).

**Figure S4.**
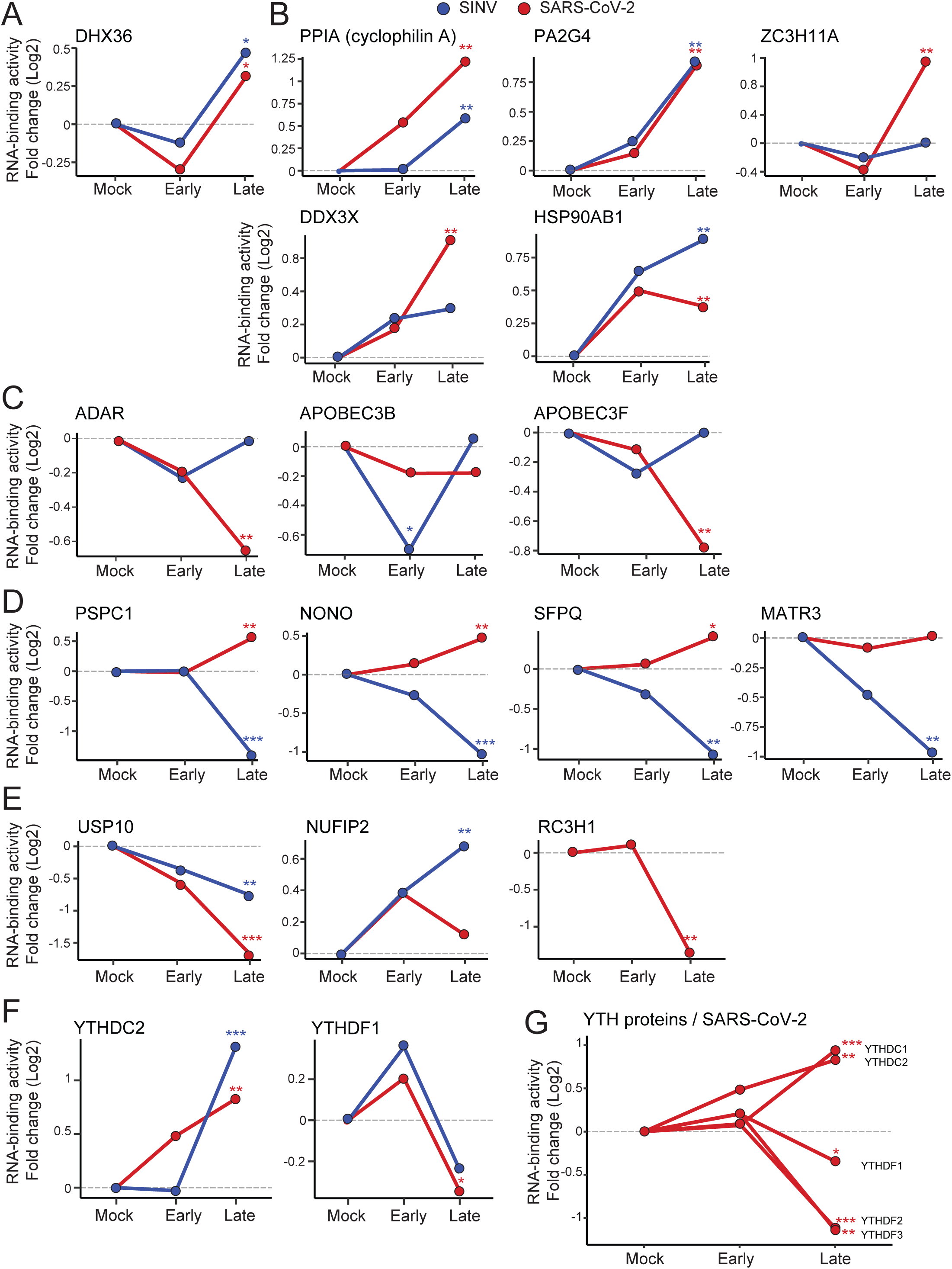
Comparison of the RBP responses to SARS-CoV-2 and SINV infection. A-F) Line plots showing the protein intensity ratio between early/mock and late/mock samples from the SARS-coV-2 (red) and SINV (blue) cRIC experiment for selected proteins. Early was defined as 8 hpi for SARS-CoV-2 and 4hpi for SINV. Late was defined as 24 hpi for SARS-CoV-2 and 18 hpi for SINV. G) Line plot showing the protein intensity ratio between 8hpi/mock and 24hpi/mock for all the YTH m6A readers detected in the cRIC experiment *, FDR < 20%; **, FDR < 10% and *** FDR < 1%.

**Figure S5.**
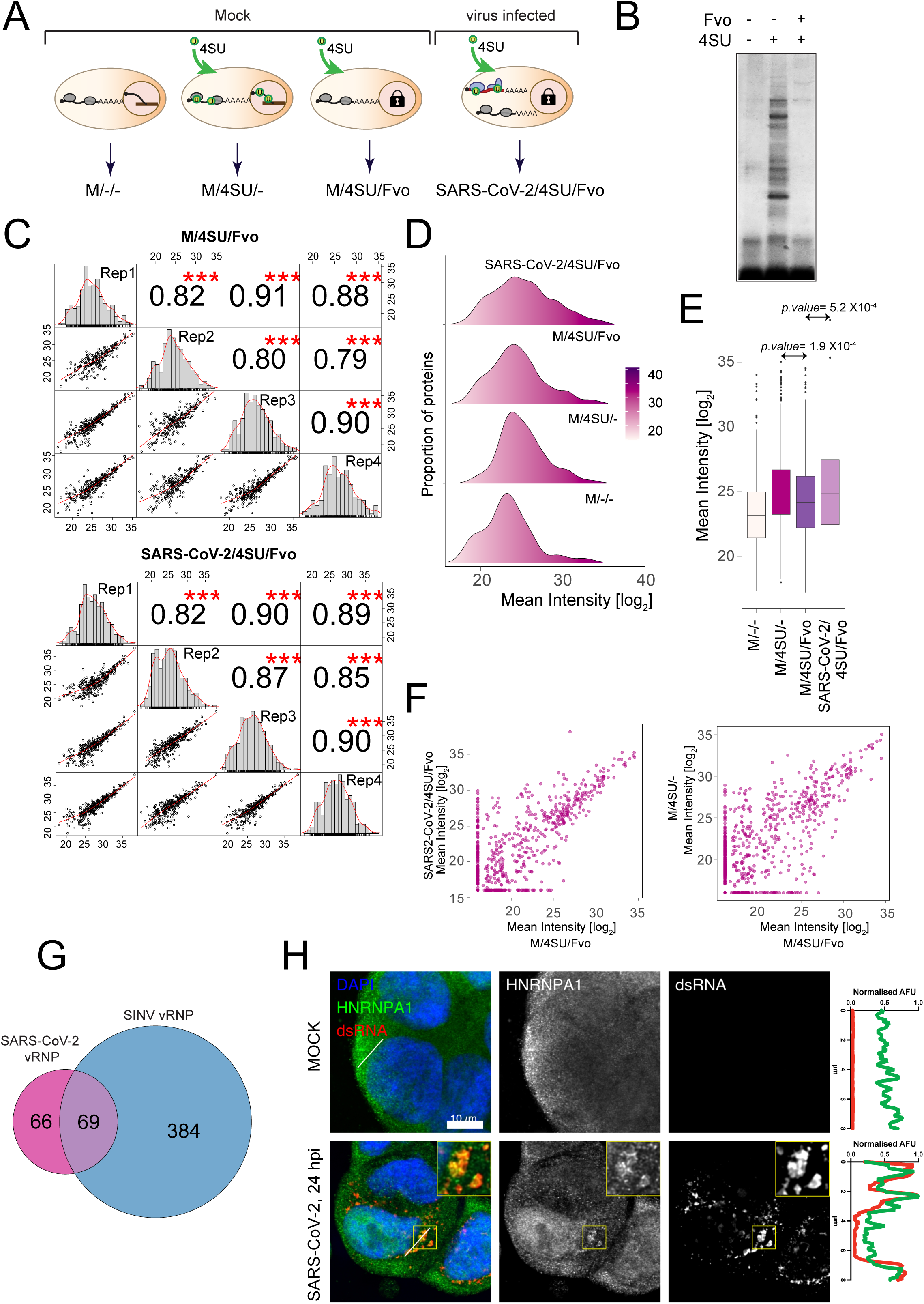
vRIC analysis of SARS-CoV-2 infected cells. A) Schematic representation of all the controls used in the vRIC experiment. B) Silver staining analysis of the inhibitory effect of Fvo in the incorporation of 4SU into poly(A) RNA in Hek293 cells. Uninfected cells were treated with or without fvo and with or without 4SU and irradiated with 365 nm UV light. RBPs bound to poly(A) RNA were isolated by RIC and eluates analysed by silver staining. Fvo strongly reduces the purification of RBPs by oligo(dT) capture suggesting lack of incorporation of 4SU into nascent RNAs. C) Scatter plots comparing protein intensity correlation between vRIC replicates for each condition. Pearson correlation is indicated. ***, p<0.001. D) Kernel density plot for the different vRIC samples showing the distribution of mean protein intensities. E) Protein intensity distribution in samples and controls of the vRIC experiment. P value is estimated by Welch’s t-test F) Correlation of protein intensity in the vRIC experiment when comparing infected vs uninfected cells and uninfected cells treated or not with Fvo. G) Venn diagram showing the overlapping between SARS-CoV-2 and SINV vRIC datasets. H) Immunofluorescence analysis using antibodies against HNRNPA1 and dsRNA in uninfected and infected cells. Right plot shows the distribution of fluorescence intensity in the green and red channels across the lines depicted in the image. AFU, arbitrary fluorescence units.

**Figure S6.**
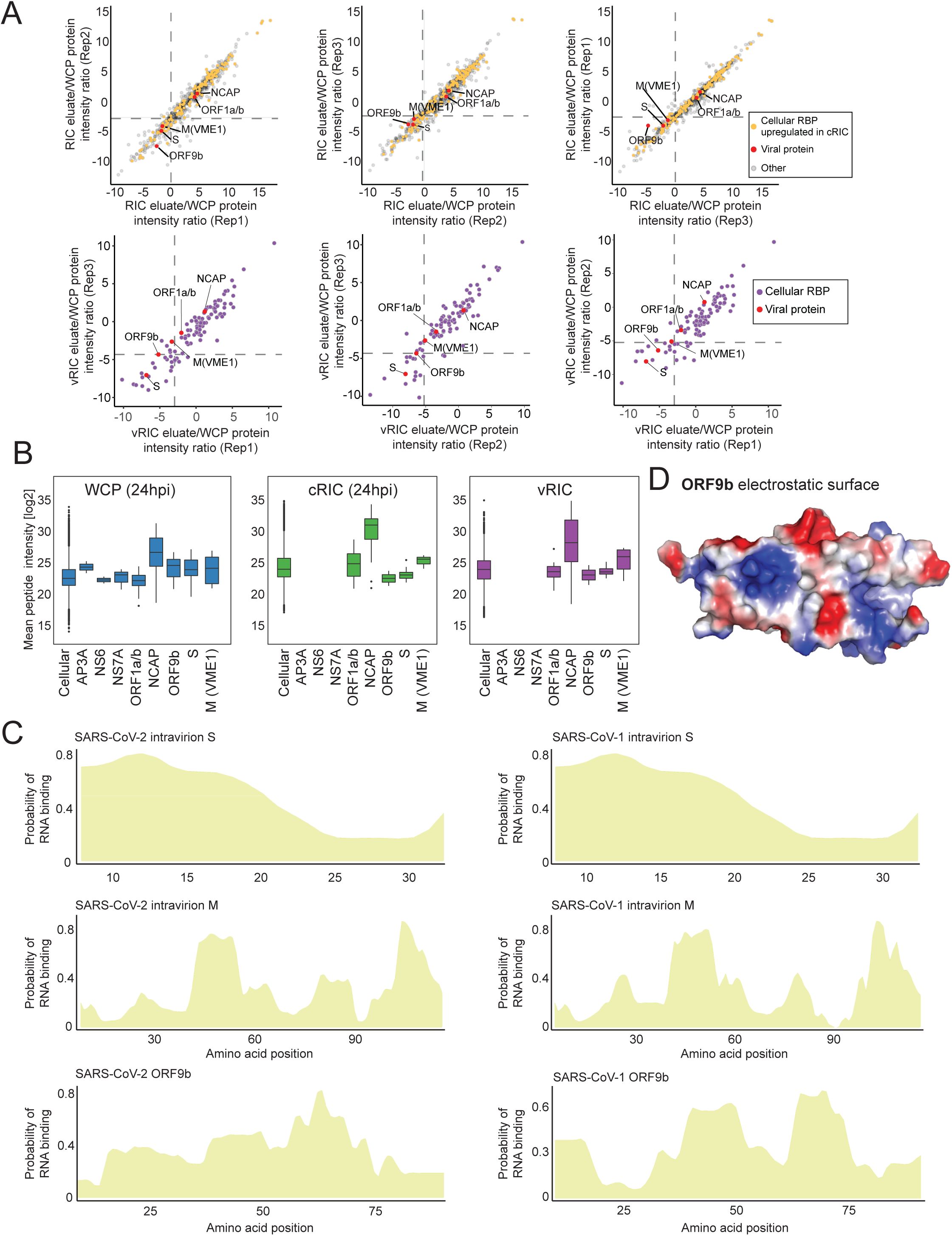
Analysis of SARS-CoV-2 proteins that interact with RNA. A) Scatter plot showing the correlation between replicates of the protein intensity ratio between cRIC and WCP (upper panels) or vRIC and WCP (bottom panels). B) Peptide intensity distribution for all the viral proteins in WCP, cRIC or vRIC at 24hpi. C) Prediction of putative RNA-binding sites within the SARS-CoV-2 (left) and SARS-CoV-1 intravirion part of S (upper panels) and M (middle panels) or full length ORF9B (bottom panels). Prediction was made with RBDetect, which employs shrinkage discriminant analysis in the positive and negative examples within the RBDmap dataset to predict RNA-binding sites based on sequence similarities with human RBPs. D) Visualisation of the electrostatic surface of ORF9b using an available 3D structure (PDB ID: 6z4u). In blue are displayed the positively charged surfaces, while the negatively charged ones are shown in red.

**Figure S7.**
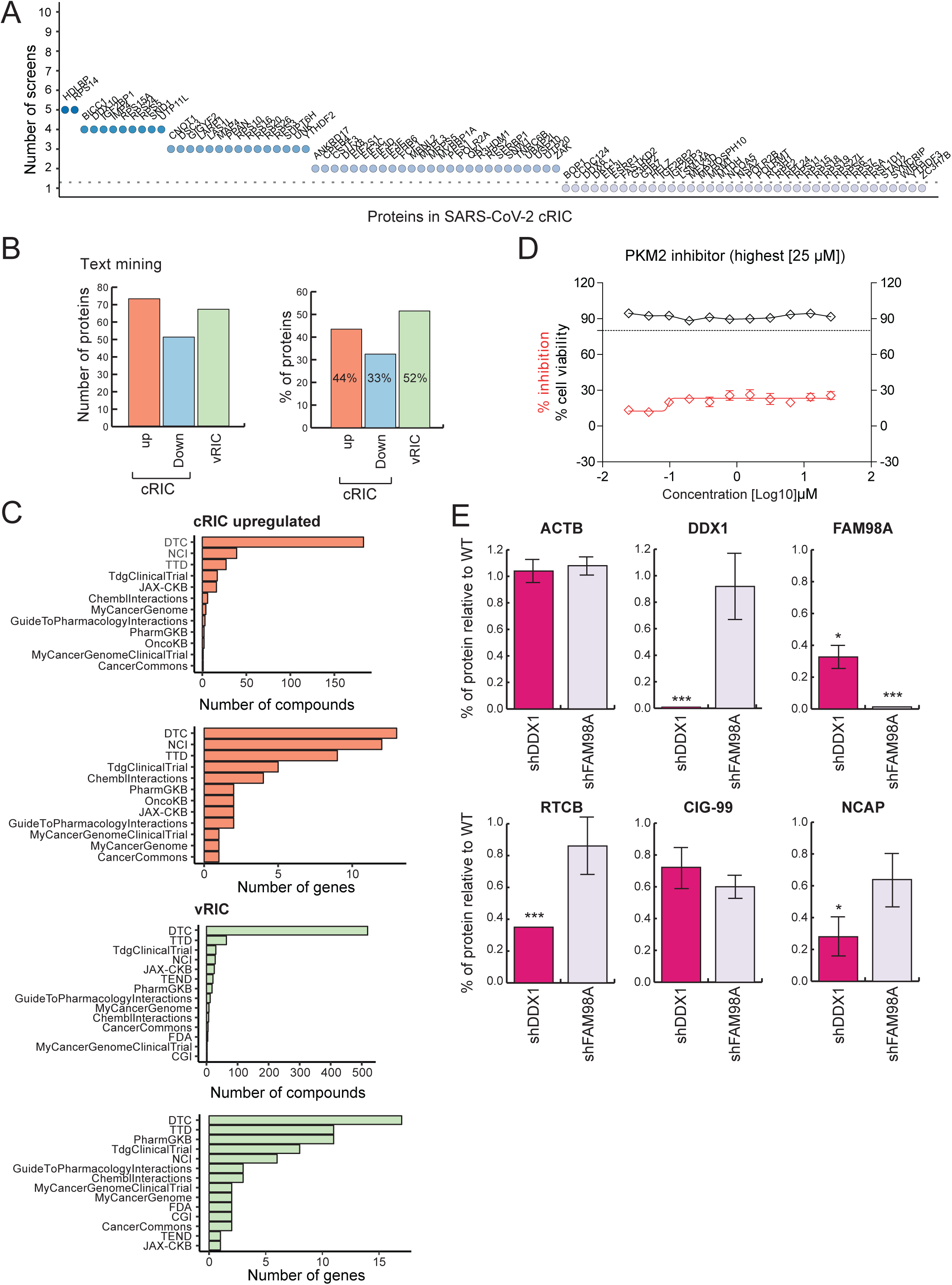
Functional implications of RBPs in SARS-CoV-2 infection. A) Proteins with identified phenotypes in genome-wide screens using viruses. Proteins dowregulated in the cRIC experiment are displayed along the x axis, while y axis indicates the number of screens in which the protein has caused a phenotype in infection. B) Proportion of proteins within the cRIC and vRIC datasets that have been linked to infection using Pudmed automatized analysis. C) Comparison of RBPs upregulated by cRIC or/and present in the vRIC dataset to drug databases. D) Effect of PKM2 inhibitor on SARS-Cov-2 infection. Red line indicates the effects in infection measured by protein ELISA at each drug dose. Black line shows cell viability at each drug dose. Error bars are SEM from three independent experiments. E) Effects of DDX1 and FAM98A knock down in the tRNA-LC subunits and SARS-CoV-2 NCAP. Data is normalised to wild type cells. Error bars represent standard deviation using information from three biological replicates. *, p< 0.05; **, p<0.01 and *** p<0.001.

## Notes

### Competing Interest Statement

The authors have declared no competing interest.

